# A monocyte/dendritic cell molecular signature of SARS-CoV2-related multisystem inflammatory syndrome in children (MIS-C) with severe myocarditis

**DOI:** 10.1101/2021.02.23.432486

**Authors:** Camille de Cevins, Marine Luka, Nikaïa Smith, Sonia Meynier, Aude Magérus, Francesco Carbone, Víctor García-Paredes, Laura Barnabei, Maxime Batignes, Alexandre Boullé, Marie-Claude Stolzenberg, Brieuc P. Pérot, Bruno Charbit, Tinhinane Fali, Vithura Pirabarakan, Boris Sorin, Quentin Riller, Ghaith Abdessalem, Maxime Beretta, Ludivine Grzelak, Pedro Goncalves, James P. Di Santo, Hugo Mouquet, Olivier Schwartz, Mohammed Zarhrate, Mélanie Parisot, Christine Bole-Feysot, Cécile Masson, Nicolas Cagnard, Aurélien Corneau, Camille Bruneau, Shen-Ying Zhang, Jean-Laurent Casanova, Brigitte Bader Meunier, Julien Haroche, Isabelle Melki, Mathie Lorrot, Mehdi Oualha, Florence Moulin, Damien Bonnet, Zahra Belhadjer, Marianne Leruez, Slimane Allali, Christèle Gras Leguen, Loïc de Pontual, Pediatric-Biocovid Study Group, Alain Fischer, Darragh Duffy, Fredéric Rieux- Laucat, Julie Toubiana, Mickaël M. Ménager

## Abstract

SARS-CoV-2 infection in children is generally milder than in adults, yet a proportion of cases result in hyperinflammatory conditions often including myocarditis. To better understand these cases, we applied a multi-parametric approach to the study of blood cells of 56 children hospitalized with suspicion of SARS-CoV-2 infection. The most severe forms of MIS-C (multisystem inflammatory syndrome in children related to SARS-CoV-2), that resulted in myocarditis, were characterized by elevated levels of pro-angiogenesis cytokines and several chemokines. Single-cell transcriptomic analyses identified a unique monocyte/dendritic cell gene signature that correlated with the occurrence of severe myocarditis, characterized by sustained NF-κB activity, TNF-α signaling, associated with decreased gene expression of NF-κB inhibitors. We also found a weak response to type-I and type-II interferons, hyperinflammation and response to oxidative stress related to increased HIF-1α and VEGF signaling. These results provide potential for a better understanding of disease pathophysiology.

## Introduction

In adults, critical forms of COVID-19 are typically characterized by severe pneumonia and acute respiratory distress syndrome (Wiersinga et al., 2020). In children, symptomatic COVID-19 occurs much less frequently and is milder than in adults, with multifactorial reasons for these differences (Brodin, 2020; Castagnoli et al., 2020; Gudbjartsson et al., 2020; Levy et al., 2020; Tagarro et al., 2020). However, in regions with high incidence of SARS-CoV-2 infection, some children have presented a postacute hyperinflammatory illness (Datta et al., 2020). In these cases, diagnostic evidence of recent SARS-CoV-2 infection has been consistently reported (Abrams et al., 2020; Jones et al., 2020; Toubiana et al., 2020, 2021). This condition was named multisystem inflammatory syndrome in children (MIS-C) or alternatively PIMS-TS (Pediatric Inflammatory Multisystem Syndrome Temporally Associated with SARS-CoV-2)(Whittaker et al., 2020). MIS-C cases most often present with symptoms similar to Kawasaki Disease (KD), an hyperinflammatory illness characterized by clinical features such as strawberry-like tongue and red and dry lips, bulbar conjunctival injection, diffuse rash, swollen extremities and cervical lymphadenopathy (McCrindle et al., 2017). KD complications can develop as myocarditis or shock syndrome in a minority of cases (Kanegaye et al., 2009). KD is thought to be triggered by viral or bacterial pathogens but the precise pathophysiological mechanisms remain elusive, with one hypothesis proposing an uncontrolled superantigen-driven inflammatory immune response (Chang et al., 2014). Compared to classic KD, MIS-C occurs in patients who are older, have more often gastrointestinal symptoms, myocarditis and shock syndrome, and exhibit higher levels of inflammatory markers (Abrams et al., 2020; Datta et al., 2020; Toubiana et al., 2020, 2021).

Inflammatory features of MIS-C are in part overlapping with those of both KD and acute SARS-CoV-2 infection in children, as well as severe COVID-19 in adults (Carter et al., 2020; Consiglio et al., 2020; Datta et al., 2020; Gruber et al., 2020). Very high levels of C-reactive protein (CRP), Procalcitonin (PCT) and IL-6, might reflect a strong immunological response to a pathogenic SARS-CoV-2 superantigen (Cheng et al., 2020). Autoimmune features can also be found in MIS-C patients (Gruber et al., 2020).

In order to further decipher SARS-CoV-2-related conditions in children, we have performed a detailed multi-parametric study combining sensitive cytokine measurements, deep immune cell phenotyping and transcriptomic analyses at the single-cell level on peripheral blood mononuclear cells (PBMCs).

## Results

### Clinical description of the cohort

The study cohort consisted of 56 children hospitalized during the first peak of the SARS-CoV-2 pandemic (from the 6^th^ of April to the 30^th^ of May), and 34 healthy controls (N=26 pediatric and N=8 adults recruited before the COVID-19 pandemic) (**Figure 1**, **Table S1**). Among the 13 children with acute respiratory infection (**Table S1**, **Figure S1**), 9 had a confirmed SARS-CoV-2 infection (RT-PCR on nasopharyngeal aspiration or swab) (*Acute-inf (CoV2^+^)* group). Six out of these 9 cases had pneumonia, and one had an uncomplicated febrile seizure. One patient with a history of recent bone marrow transplantation for sickle cell disease, required intensive care support. The 4 other patients (*Acute-inf (CoV2^-^)* group) had pneumonia associated with a positive RT-PCR test for either *Mycoplasma pneumoniae* or rhinovirus/enterovirus, and negative RT-PCR for SARS-CoV-2.

**Figure 1:**
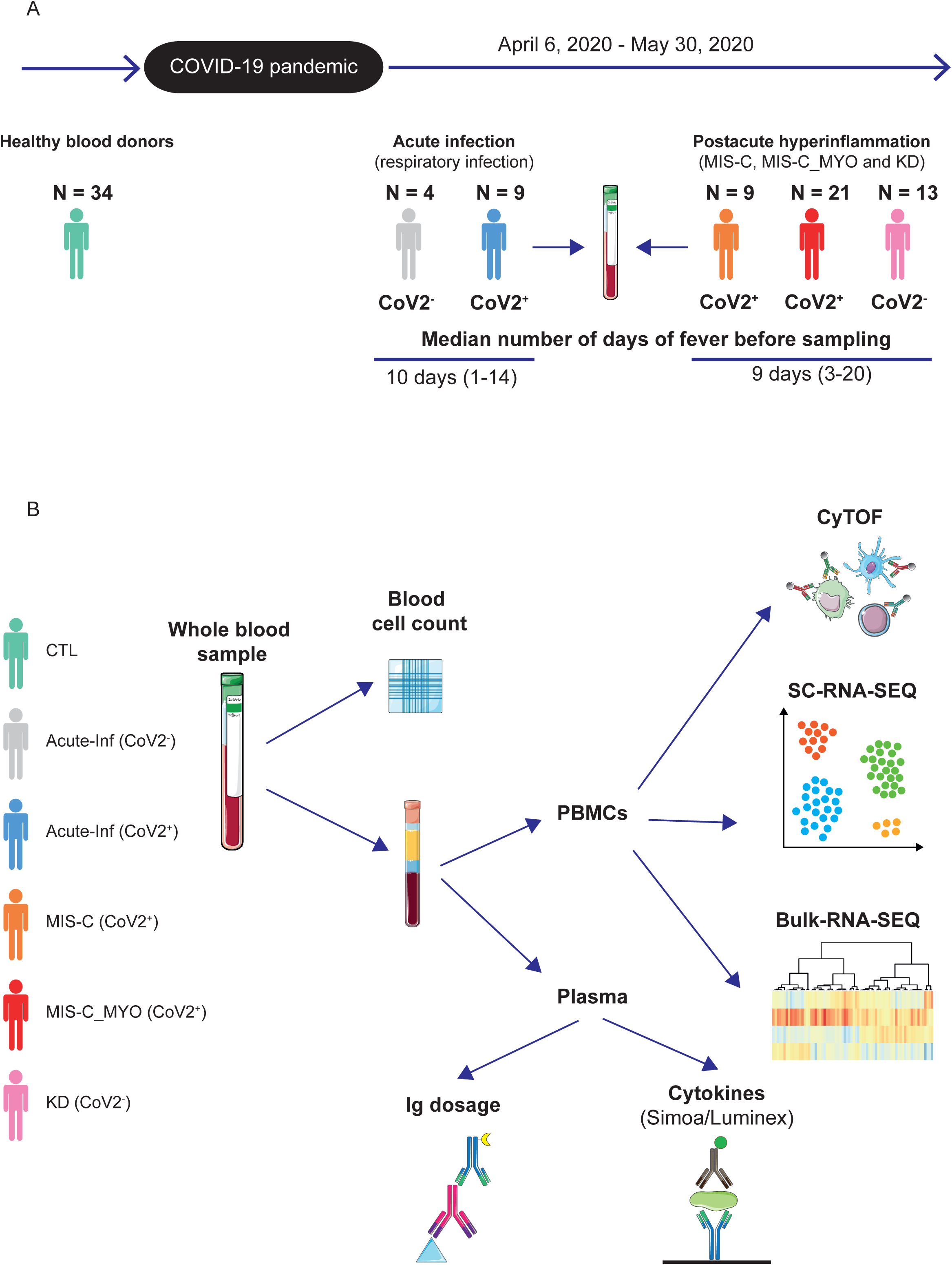
Timeline and experimental designs. **A.** Timeline depicting when the different groups of pediatric patients were enrolled. **B.** Description of the different types of analyses performed on whole blood samples, peripheral blood mononuclear cells (PBMCs) and plasma. CyTOF: Mass cytometry (Cytometry by Time Of Flight). SC-RNA-SEQ: single-cell transcriptomic followed by sequencing. Bulk-RNA-SEQ: transcriptomic bulk level sequencing. Simoa: Single molecule array, digital ELISA. Luminex: cytokine bead array assays. Ig dosage; quantification of SARS-CoV-2 specific immunoglobulins. CTL, healthy donors, in green; *Acute-inf (CoV2^-^)*, patients with acute respiratory infection but no evidence of SARS-CoV-2 infection, in gray; *Acute-inf (CoV2^+^)*, patients with acute respiratory infection and evidence of SARS-CoV-2 infection, in blue; *MIS-C (CoV2^+^)*, patients with postacute multi-inflammatory syndrome and evidence of SARS-CoV-2 infection, in orange; *MIS-C_MYO (CoV2^+^)*, patients with postacute hyperinflammatory syndrome, severe myocarditis and evidence of SARS-CoV-2 infection, in red; *KD (CoV2^-^)*, patients with postacute hyperinflammatory syndrome, no evidence of SARS-CoV-2 infection, but criteria for Kawasaki Disease (KD), in pink. Illustrations were obtained from Servier Medical Art, licensed under a Creative Common Attribution 3.0 Generic License. http://smart.servier.com/.

Forty-three children displayed features of postacute hyperinflammatory illness (**Figure S1**, **Table S1**). SARS-CoV-2 infection status of all samples was confirmed by specific antibody determination (both IgG and IgA) in the plasma, using ELISA and flow cytometry-based technics as described in the methods (**Figure S2A**). Most (n=30) had a confirmed SARS-CoV-2 infection (with 14 also positive for concomitant nasopharyngeal RT-PCR testing) and were therefore considered as cases of MIS-C (*MIS-C (CoV2^+^)* group); all 30 cases of MIS-C presented clinical features of KD, 14 of them fulfilled clinical criteria for a complete form of KD according to the American Heart Association (McCrindle et al., 2017). Of note, 21/30 cases had severe myocarditis (i.e. with elevated high-sensitivity cardiac troponin I and/or regional wall motion abnormalities on echocardiography, and clinical signs of circulatory failure requiring intensive care support; *MIS-C_MYO (CoV2^+^)*). Thirteen tested negative for SARS-CoV-2 and fulfilled clinical criteria for complete (n=6) or incomplete (n=7) Kawasaki disease (KD), and were therefore considered to have KD-like illness (*KD (CoV2^-^)* group) (**Figure S1**, **Table S1**). Clinical and biological characteristics, at time of disease activity and before treatment, or within 24 hours of treatment onset, are presented in **Table S1.** MIS-C cases had low lymphocyte counts and those with severe myocarditis had in addition abnormally increased neutrophil counts as compared to other groups, along with high levels of CRP, PCT, serum alanine transaminases (ALT) and ferritin (**Table S1**). All cases responded favorably to intravenous immunoglobulin injections (IVIG), some (N=12, Table S1) in combination with glucocorticosteroids, received before sampling. Multi-parametric analyses were performed at a median fever persistence of 9-10 days (**Figures 1A, B**).

### Elevated inflammatory cytokine levels in pediatric acute infection and postacute hyperinflammatory conditions

We investigated plasma cytokine and chemokine levels in all patients, by Luminex and Simoa assays. Hierarchical clustering analysis and stratification by patient groups revealed overall elevated levels of immune and inflammatory markers, with 40/46 measured proteins significantly elevated (q<0.05) as compared to healthy controls (**Figure 2A**; global heat map). Twelve cytokines were found to be elevated in all groups of patients as compared to healthy controls **(Figure S2B).** High IL-8 and CXCL1 (**Figure S2C**) were more specific to children with acute infection. Cytokine levels did not significantly differ between children with acute infection with or without evidence of SARS-CoV-2 infection (**Figures S2D-F**). IFNγ, IFNα2, IL-17A, TNF-α, IL-10, Granzyme B, were higher in children with postacute hyperinflammation (*MIS-C (CoV2^+^)*, *MIS-C_MYO (CoV2^+^)*, and *KD (CoV2^-^)* groups), as compared to pediatric healthy donors (CTL) and patients with acute infections (*Acute-inf (CoV2^+^)* and *Acute-inf (CoV2^-^)* (**Figure 2A**, top cluster **and 2B).** A slightly higher expression of IFNα2 and IL-17A was found in MIS-C without myocarditis (*MIS-C (CoV-2*^+^) patients). (**Figure 2C**). In contrast, 11 cytokines and chemokines (CSF2, CCL2, IL-6, CXCL10, FLT3L, VEGF, TGF-A, IL-1RA, PD-L1, CX3CL1, TGF-B1) were higher in MIS-C with severe myocarditis (*MIS-C_MYO (CoV-2*^+^)) (**Figures 2 C, D**). Of note, 8 of them are known to be associated with TNF-α signaling. They are involved in propagation of inflammation (IL-6, IL-15), angiogenesis and vascular homeostasis (VEGF and TGF cytokines) and activation and chemotaxis of myeloid cells (CCL2, CX3CL1, CXCL10) (**Figure 2D**) (Holbrook et al., 2019; Varfolomeev and Ashkenazi, 2004). An increased level of CCL19, CCL20, CCL3 (cell migration and chemotaxis) and IL-1 agonist/antagonist (IL-1β, IL-1RA) were also observed, as well as increased soluble PD-L1 (**Figure S2G**). Another noticeably elevated cytokine was CSF2, known to be involved in myeloid cell differentiation and migration (**Figure 2D**).

**Figure 2:**
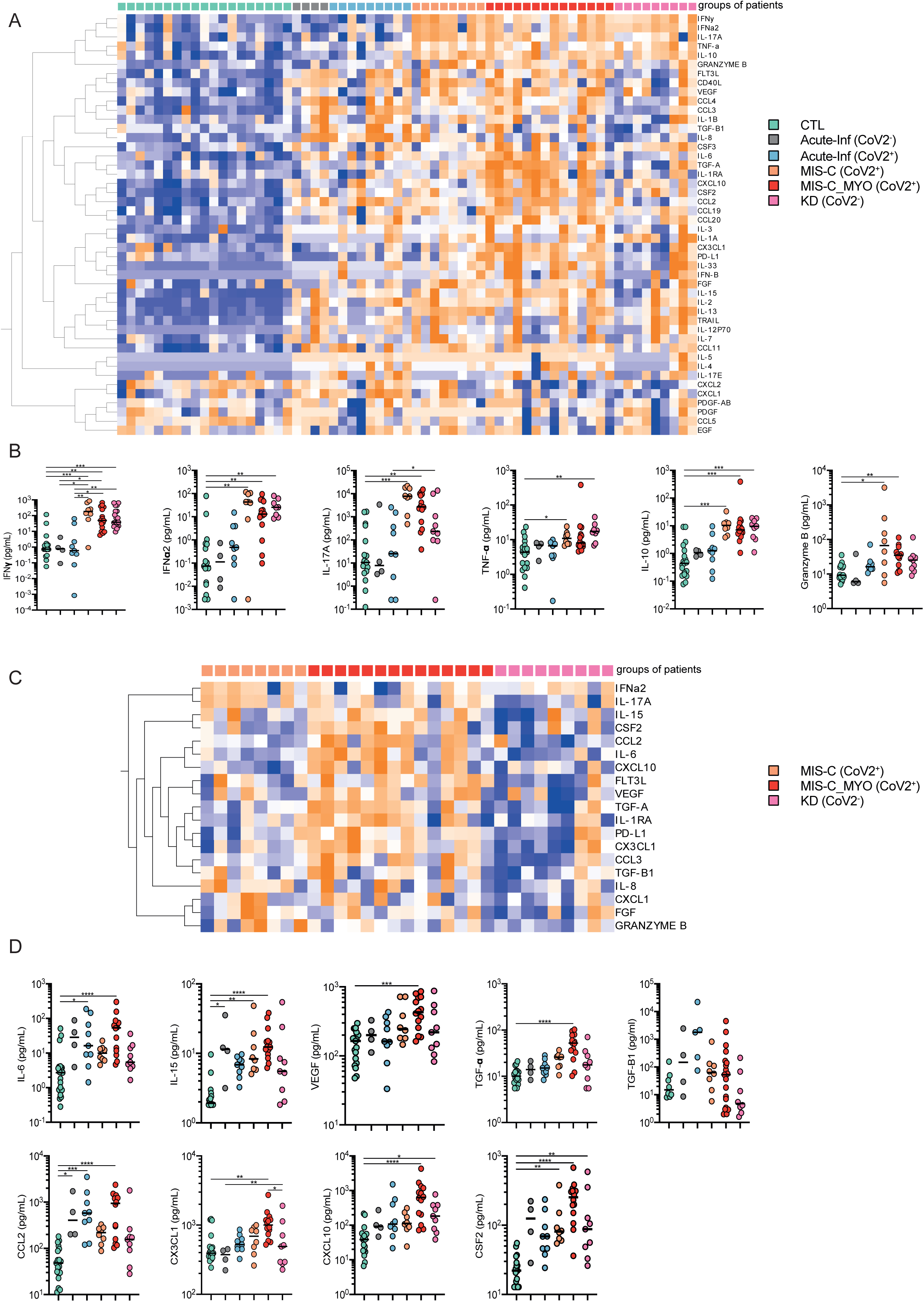
Analyses of cytokine plasma levels. **A.** Heatmap of the cytokines differentially secreted between the different clinical groups: CTL, green; *Acute-inf (CoV2^-^)*, gray; *Acute-inf (CoV2^+^)*, blue; *MIS-C (CoV2^+^)*, orange; *MIS-C_MYO (CoV2^+^)*, red; *KD (CoV2^-^)*, pink). On the x axis, blood donors are organized by groups and on the y axis, cytokines are displayed following hierarchical clustering. Cytokines were expressed as pg/ml and log transformed with blue to orange colors representing lower to higher expression respectively. **B.** Dot plots of cytokines elevated in postacute hyper inflammatory groups (*MIS-C (CoV2^+^)*, *MIS-C_MYO (CoV2^+^)* and *KD (CoV2^-^)*), as compared to early infection groups (*Acute-inf (CoV2^+^)*, *Acute-inf (CoV2^-^)*) and healthy blood donors (CTL). **C.** Heatmap of the cytokines differentially expressed among the patients with postacute hyper inflammation: *MIS-C (CoV2^+^)*, *MIS-C_MYO (CoV2^+^)* versus *KD (CoV2^-^)*). **D.** Dot plots of cytokines with a higher expression in the *MIS-C_MYO (CoV2^+^)*. **B & D**. ρ-values are calculated by Kruskal-Wallis test for multiple comparisons, followed by a post-hoc Dunn’s test. * (ρ < 0.05), **(ρ < 0.01), ***(ρ < 0.001)

Altogether, high inflammatory cytokine levels were detected in both acute infection and postacute inflammatory cases. The strongest inflammatory profile was observed in MIS-C with severe myocarditis (*MIS-C_MYO (CoV2*^+^)) and remarkably, a similar profile was observed when comparing MIS-C without myocarditis (*MIS-C (CoV2*^+^)) and KD-like illness unrelated to SARS-CoV-2 (*KD (CoV2^-^)*). Of note, the inflammatory profile was much reduced in intensity in MIS-C cases with myocarditis under combined glucocorticosteroid and IVIG treatment, as compared to IVIG alone (**Figure S2H**).

### Low monocyte and dendritic cell frequencies in patients with postacute hyperinflammatory illness

To better decipher the blood immune cell composition of each group, PBMCs were analyzed by CyTOF mass spectrometry and by single-cell analyses at the transcriptomic level (SC-RNA-SEQ) (**Figure 1**). Clustering analyses of the data obtained from CyTOF and SC-RNA-SEQ revealed consistent results, with most of the alterations observed in clusters composed of monocytes or dendritic cells (**Figures 3A, B; S3 A-D**). The most drastic changes were a decrease in conventional Dendritic Cells (cDCs) and plasmacytoid Dendritic Cells (pDCs) in patients with a postacute hyperinflammatory illness ((*MIS-C_MYO (CoV2*^+^*), MIS-C (CoV2*^+^) and *KD (CoV2^-^)*). As previously reported (Gruber et al., 2020), we also observed a trend towards a decrease in monocyte clusters in children with postacute hyperinflammatory illness, that was found independent of SARS-CoV-2 status (**Figures 3A, B; S3 A-D**). Of note, some heterogeneity was observed in the proportions of non-classical monocytes in *Acute-Inf (CoV2^+^)* cases. We observed additional heterogeneity in the proportions of classical and intermediate monocytes in patients with severe myocarditis (*MIS-C_MYO (CoV2*^+^) (**Figures 3A, B**), but there was no correlation with clinical data (**Table S1**), nor cytokine/chemokine measurements (**Figures 2 and S2**). Additional modifications were detected in patients with acute SARS-CoV-infection (*Acute-inf (CoV2^+^)* cases), consisting of a decrease of MAIT cells and an excess of naïve and central memory CD4^+^ T cells (**Figures 3 A, B, S3 C, D**).

**Figure 3:**
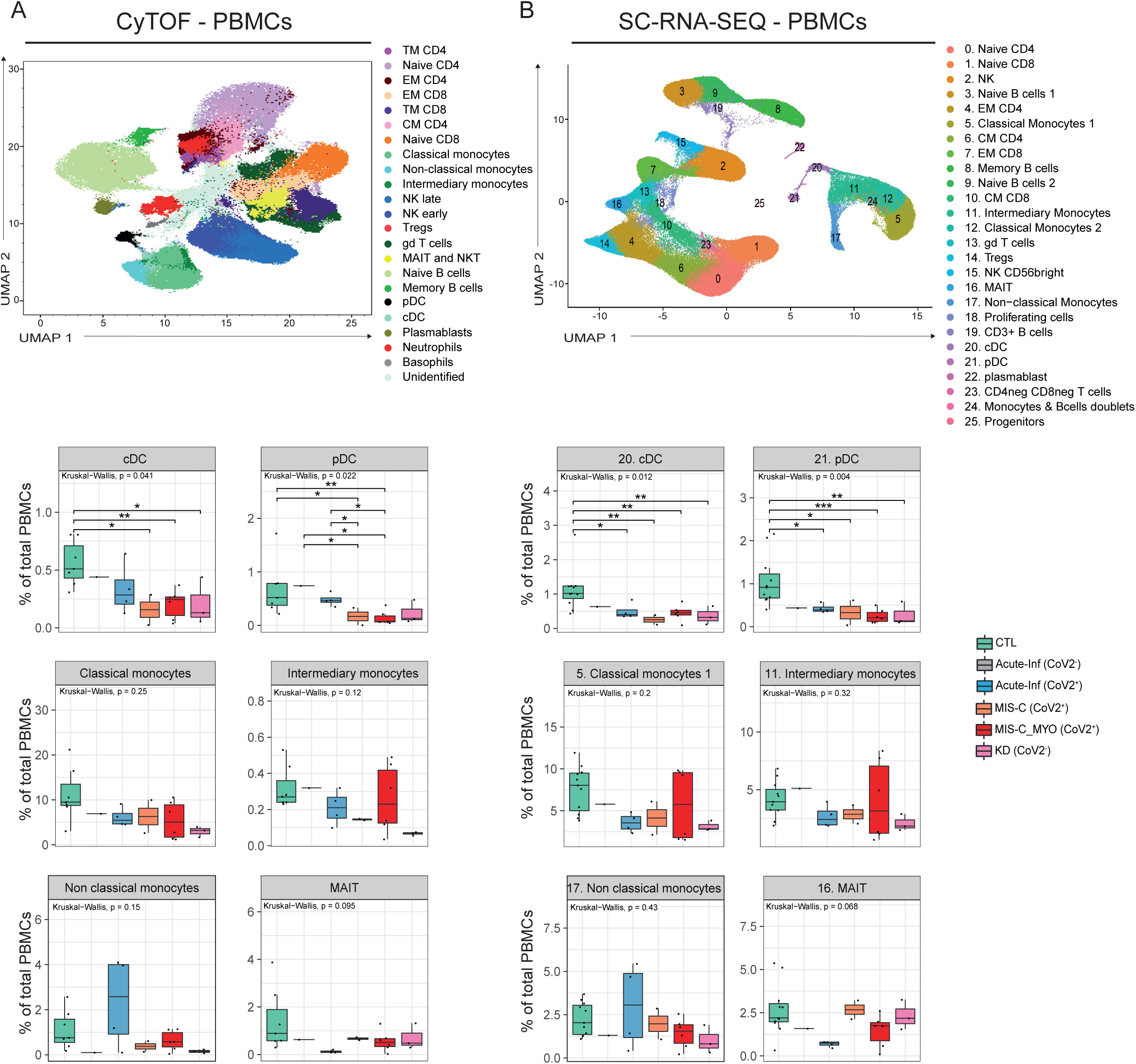
CyTOF and SC-RNA-SEQ characterization of peripheral blood immune cell (PBMCs) distribution. **A.** Upper panel: UMAP of 1,150,000 single cells from PBMCs of 7 CTL, 1 *Acute-inf (CoV2^-^)*, 4 *Acute-inf (CoV2^+^)*, 2 *MIS-C (CoV2^+^)*, 6 *MIS-C_MYO (CoV2^+^)* and 3 *KD (CoV2^-^)* donors, following analyses by CyTOF and displayed as 23 clusters identified using the individual expression of 29 proteins, as described in Figure S3A. Bottom panel: boxplots of clusters with significant differences observed between groups. **B.** Upper panel: UMAP of 152,201 single cells following extraction from PBMCs (9 CTL, 1 *Acute-inf (CoV2^-^)*, 4 *Acute-inf (CoV2^+^)*, 2 *MIS-C (CoV2^+^)*, 6 *MIS-C_MYO (CoV2^+^)*, and 3 *KD (CoV2^-^)*) and processed by SC-RNA-SEQ. A resolution of 0.8 allows to segregate cells into 26 clusters identified based on the expression of several markers and gene signatures, as shown in Figure S4B. Bottom panel: Boxplots of clusters with significant differences. **A & B.** (CTL, green; *Acute-inf (CoV2^-^)*, gray; *Acute-inf (CoV2^+^)*, blue; *MIS-C (CoV2^+^)*, orange; *MIS-C_MYO (CoV2^+^)*, red; *KD (CoV2^-^)*, pink). In the boxplots, each dot represents a sample. Boxes range from the 25th to the 75th percentiles. The upper and lower whiskers extend from the box to the largest and smallest values respectively. Any samples with a value at most x1.5 the inter-quartile range of the hinge is considered an outlier and plotted individually. ρ values are calculated by Kruskal-Wallis test for multiple comparisons, followed by a post hoc Dunn’s test. * (ρ < 0.05), **(ρ < 0.01), ***(ρ < 0.001).

### Overexpression of inflammatory pathways, NF-κB signaling, and metabolic changes related to hypoxia in acute infection and postacute hyperinflammatory conditions

To gain further insight into the mechanisms behind the different forms of SARS-CoV-2 infection in children, we assessed pathways modulated in each group by looking at the differentially expressed genes, obtained from the SC-RNA-SEQ dataset. In patients with acute infection (*Acute-inf (CoV2^+^) and Acute-inf (CoV2^-^)*), the numbers of differentially expressed genes were homogeneously distributed among monocytes/DCs, T and B cells (**Figures 4A and S4A).** Pathway enrichment analyses (Ingenuity Pathway Analyses; IPA, QIAGEN Inc.) revealed a decrease in oxidative phosphorylation, coupled with an increase of HMGB1 signaling, HIF-1α signaling, and hypoxia signaling. Production of nitric oxide was observed in both groups of acute infections, independently of SARS-CoV-2 infection, as compared to healthy controls (**Figure S4B**). These observations suggest a metabolic switch potentially driven by hypoxic conditions. NF-κB signaling, VEGF signaling and inflammatory pathways (type-I and type-II IFNs, IL-1, IL-6, and IL-17 signaling pathways) were also found to be overrepresented in both groups of patients (**Figure S4B**).

**Figure 4:**
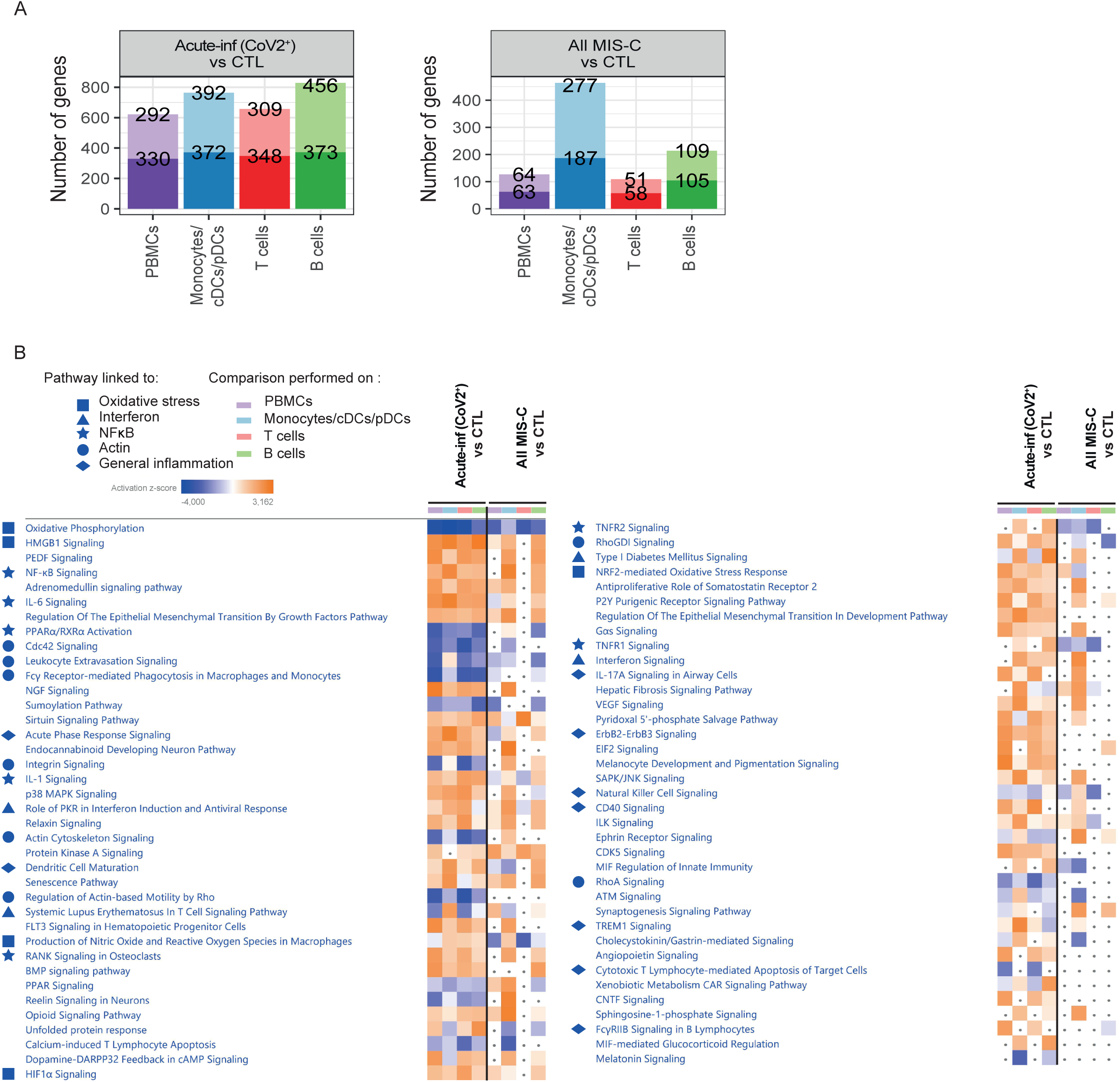
Genes and pathways differentially regulated in acute infection and postacute hyper inflammation following SARS-CoV-2 infection. **A & B.** Bar charts of the number of up- and downregulated genes in *Acute-inf (CoV2^+^)* (A) and All MIS-C (*MIS-C (CoV2^+^)* and *MIS-C_MYO (CoV2^+^)*) (B), compared to CTL, in PBMCs, monocytes/DCs, T and B cells clusters obtained following SC-RNA-SEQ experiments as displayed in Figure 3B. PBMCs represent all clusters; monocytes/DCs cells, clusters 5, 11, 12, 17, 20, 21, 24; T cells, clusters 0, 1, 2, 4, 6, 7, 10, 13, 14, 15, 16, 18 and 23; and B cells, clusters 3, 8, 9, 19, and 22. The top value on the light-colored bars represent the upregulated genes and the bottom dark represent the downregulated genes. **B.** Heatmap of the canonical pathways, enriched in the differentially expressed genes (DEG) from the comparisons performed in A and B in PBMCs, monocytes/DCs, T and B cells, obtained by using Ingenuity Pathways Analysis (IPA) analyses. Left panel, part 1 and right panel part 2. Symbols are used in front of the pathways to represent pathways belonging to the same functional groups. Pathways with an absolute z-score ≤ 2 or adjusted p-value > 0.05 in all conditions were filtered out. Z-score > 2 means that a function is significantly increased (orange) whereas a Z-score < −2 indicates a significantly decreased function (blue). Grey dots indicate non-significant pathways (ρ > 0.05).

Interestingly, alterations in the same pathways were also identified in all cases of children with SARS-CoV-2-related postacute illnesses (*All MIS-C (CoV2^+^)*: *MIS-C_MYO (CoV2*^+^) and *MIS-C (CoV2*^+^)). However, in these cases, alterations were mostly restricted to monocytes and dendritic cells (**Figures 4 A, B**). Comparisons of genes differentially expressed between children with postacute hyperinflammatory illness with or without evidence of SARS-CoV-2 infection (*All MIS-C (CoV2^+^)* versus *KD (CoV2^-^)*), did not reveal significant differences with the exception of type-I and type-II interferon signaling (**Figures S4 C, D)**.

The NF-κB signaling pathway was identified to be activated in monocytes and DCs of all patients with acute infection and postacute hyperinflammatory illness, independently of SARS-CoV-2 infection (**Figure 5A**). While monocytes and DCs of patients with acute infection (*Acute-inf (CoV2^+^)* and *Acute-inf (CoV2^-^)*) highly expressed genes of the NF-κB complex (*REL*, *RELA*, *RELB*, *NFKB1*, *NFKB2*; **Figures 5B, D**), monocytes and DCs from all MIS-C patients (*MIS-C_MYO (CoV2*^+^) and *MIS-C (CoV2*^+^)) exhibited a strong decrease in the expression of NF-κB inhibitors, such as *TNFAIP3* (A20), *TNFAIP2*, *NFKBIA, NFKBID, NFKBIE* and *NFKBIZ* (**Figures 5C, D**). Strikingly, this decrease in the expression of NF-κB inhibitors appeared to be specific to the monocytes and DCs of MIS-C patients with severe myocarditis (*MIS-C_MYO (CoV2*^+^)) (**Figures 5 E, S5 A-D**).

**Figure 5:**
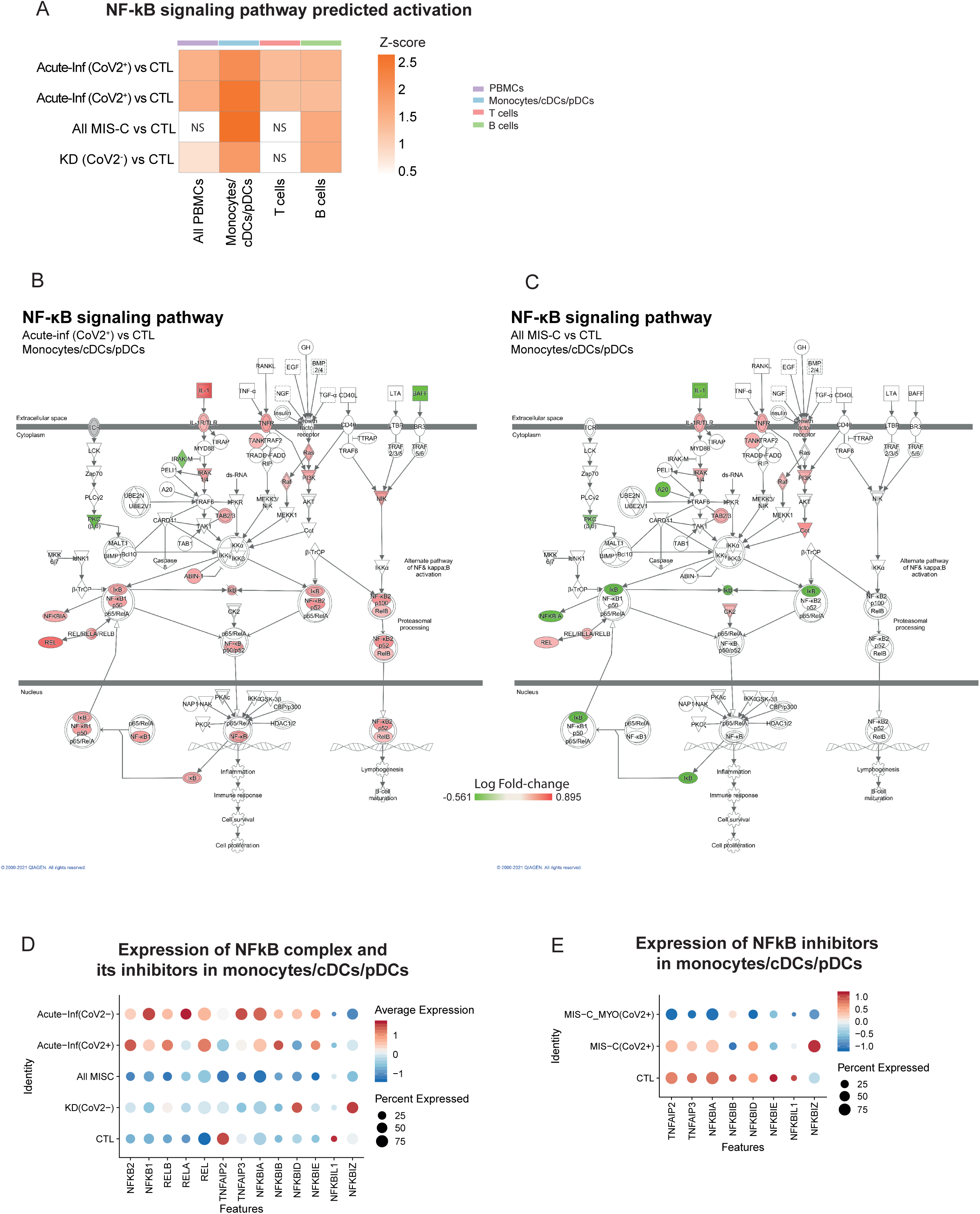
NF-κB signaling pathway activation in acute infection and postacute hyperinflammatory illness. **A.** Heatmap of the activation of NF-κB signaling pathway, as predicted by IPA, in each disease group compared to controls. Color scale represents the z-score of the prediction. The higher the score, the more activated the NF-κB signaling pathway. “NS” indicates non-significant comparisons. **B & C.** NF-κB signaling pathway from the IPA software, displayed in the monocytes/DCs cells in *Acute-inf (CoV2^+^)* versus CTL (B) or *All MIS-C (CoV2^+^)* versus CTL (C). Changes in gene expression are represented by the log fold change (increase in red and decreased in green). **D.** Dot plot of the expression in monocytes/DCs cells of the positive (from *NFKB2 to REL*) and negative regulators (from *TNFAIP2* to *NFKBIZ*) of NF-κB complex in all disease groups. **E.** Dot plot of the expression in monocytes/DCs of the negative regulators of NF-κB complex in the MIS-C groups.

In conclusion, pathways dysregulated in acute infection or postacute hyperinflammatory illness, reflected an inflammatory status based on NF-κB signaling combined with changes in metabolism driven by a hypoxic environment. In acute respiratory disease, changes in gene expression reflected involvement of all PBMCs, whereas in postacute hyperinflammatory illnesses, monocytes/DCs were the most affected populations. These results further emphasize the role of monocytes/DCs populations in MIS-C.

### Molecular pathways specific to MIS-C with severe myocarditis

To gain further insight into the inflammatory phenotype of monocytes and DCs from MIS-C patients, we compared gene expression between MIS-C patients with or without severe myocarditis and healthy controls. Most gene alterations happened in monocytes/DCs **(Figure S6A and S6C)**, which lead us to focus on these populations for the following analyses. Pathway enrichment performed both on IPA and EnrichR (Chen et al., 2013; Kuleshov et al., 2016) highlighted the modulation of type-I and type-II interferon signaling pathways, with the upregulation of several interferon stimulated genes (ISGs) (*JAK2*, *STAT1*, *STAT2*, *IFITM1, IFITM2, IFI35, IFIT1, IFIT3, MX1, IRF1*) only in the monocytes and DCs of MIS-C patients without myocarditis (**Figures 6 A, B, S6 B, D**). However, both groups of MIS-C patients showed elevated plasma IFN-α2 and IFNγ proteins (**Figures 2A, B**). Gene expression downregulation in monocytes and DCs of MIS-C patients with severe myocarditis, included most of the MHC class II genes suggesting a decrease in antigen processing and presentation pathways (**Figures 6 C and S6D**).

**Figure 6:**
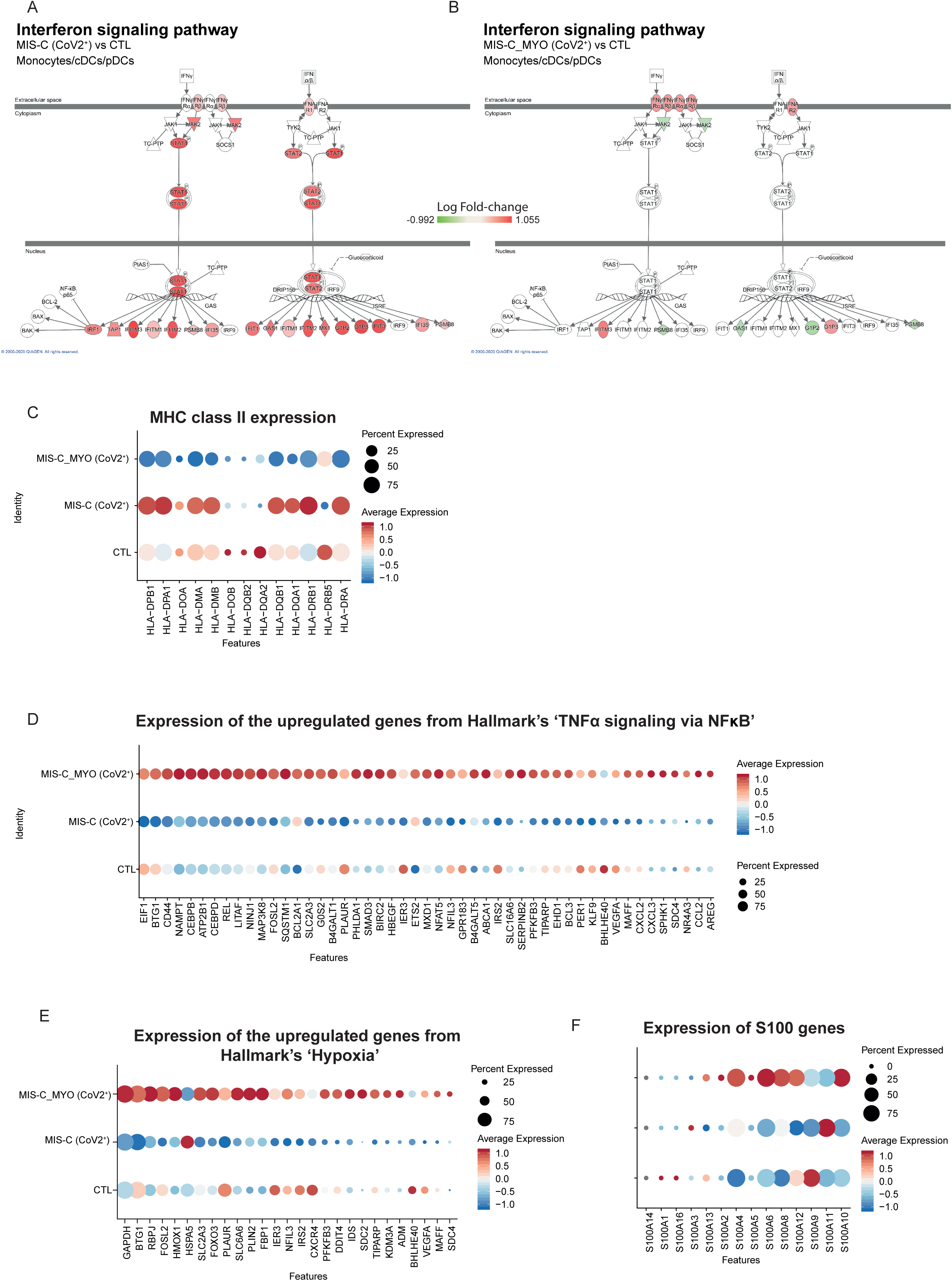
Key pathways and genes deregulated in monocytes/DCs cells of MIS-C_MYO (CoV2^+^) compared to MIS-C (CoV2^+^). **A & B.** Type-I and type-II interferon signaling pathways from the IPA software. Changes in gene expression are represented by the log fold change in monocytes/DCs cells of *MIS-C (CoV2^+^)* (A) or *MIS-C_MYO (CoV2^+^)* (B) compared to CTL (red color represents an upregulation, green a downregulation and white no changes). **B.** Dot plot of the MHC class II-associated genes. **D.** Dot plot of the 49 genes from the TNF-alpha Signaling via NF-κB pathway (pathway enrichment analysis by MSigDB Hallmark 2020 (Figure S6E)). **E.** Dot plot of the 27 genes from the Hypoxia pathway (pathway enrichment analysis by MSigDB Hallmark 2020 (Figure S6E)). **F.** Dot plot of the calcium binding genes of the S100 family in CTL, *MIS-C (CoV2^+^)* and *MIS-C_MYO (CoV2^+^)* **C - F.** The expression is represented by the centered scale expression of each gene. The “percent expressed” represents the percentage of cells that express the gene.

Transcription factor prediction in EnrichR revealed overexpression of targets of the NF-κB complex in MIS-C patients with severe myocarditis as compared to MIS-C without myocarditis (**Figure S6E**). This activation of NF-κB signaling in MIS-C patients with severe myocarditis was found to be associated with the strong downregulation of NF-κB inhibitors as shown in **Figure 5E**.

A strong overexpression of genes belonging to TNF-α signaling, as well as inflammatory responses, hypoxia and response to oxidative stress (*HIF1A*, *HMOX1*, *HMBG1, etc.)* was found in cases with severe myocarditis (**Figures 6D, E and S6E**). This was associated with a downregulation of genes linked with oxidative phosphorylation, nitric oxide production and iNOS signaling (**Figure S6B**). TGF-β signaling and VEGF signaling were also found enriched in monocytes and DCs of patients with myocarditis and to a lesser magnitude in B cells (**Figures S6 B, E).** Interestingly, an increased expression of several genes encoding S100 proteins and calcium-binding cytosolic proteins, all known to serve as danger signals to regulate cell migration, homeostasis and inflammation, were noticed in the cases of severe myocarditis (**Figure S6F**) (Xia et al., 2018).

To summarize, NF-κB activation, a decreased expression of NF-κB inhibitors, TNF-α signaling, together with a hypoxic response to oxidative stress, VEGF signaling, downregulation of MHC-II genes and low type-I and type-II IFN responses characterize the monocytes and DCs of children with MIS-C and severe myocarditis.

### Identification of a molecular signature specific to MIS-C with severe myocarditis

To identify a potential clinical relevance of our study, we searched for a molecular signature that correlated with the appearance of severe myocarditis among the monocytes/DCs of children with SARS-CoV-2-related MIS-C. By using several SC-RNA-SEQ comparison strategies (**Figure 7A**), we identified 329 genes upregulated in monocytes and DCs of the MIS-C group with myocarditis (N=6) as compared to all other groups (**Figure 7A**). To validate this molecular signature, RNA from PBMCs were sequenced from an independent group of patients. A scoring system was generated, based on normalized expression represented by a Z-score, coupled with hierarchical clustering, in order to identify genes that were overexpressed in children with myocarditis (*MIS-C_MYO (CoV2^+^)* group) as compared to the other groups (**Figure S7A**). Within the 329 genes identified by SC-RNA-SEQ in monocytes and DCs of patients with severe myocarditis, expression of 116 genes were found upregulated in PBMCs from an independent group of 9 patients belonging to the *MIS-C_MYO (CoV2^+^)* group with myocarditis (**Figures 7B**). From these genes, a signature score (SignatureSCORE) was determined for each sample processed by Bulk-RNA-SEQ (**Figure 7C**). We then further developed a RankingSCORE (**Figures S7 A, B**) to identify the top genes that contributed the most to the monocytes and DCs myocarditis signature. This led to the identification of a set of 25 genes that clearly segregate patients with severe myocarditis from other MIS-C and *KD (CoV2^-^)* (**Figure 7D)**. Consistently, most of these 25 genes belong to functional pathways that were previously identified (**Figures 6 and S6**), such as inflammation, oxidative stress, TNF-α and/or NF-κB signaling, and in some cases already known markers of myocarditis or MIS-C and/or COVID-19, such as genes coding for S100 proteins (**Figures S7 C, D**).

**Figure 7:**
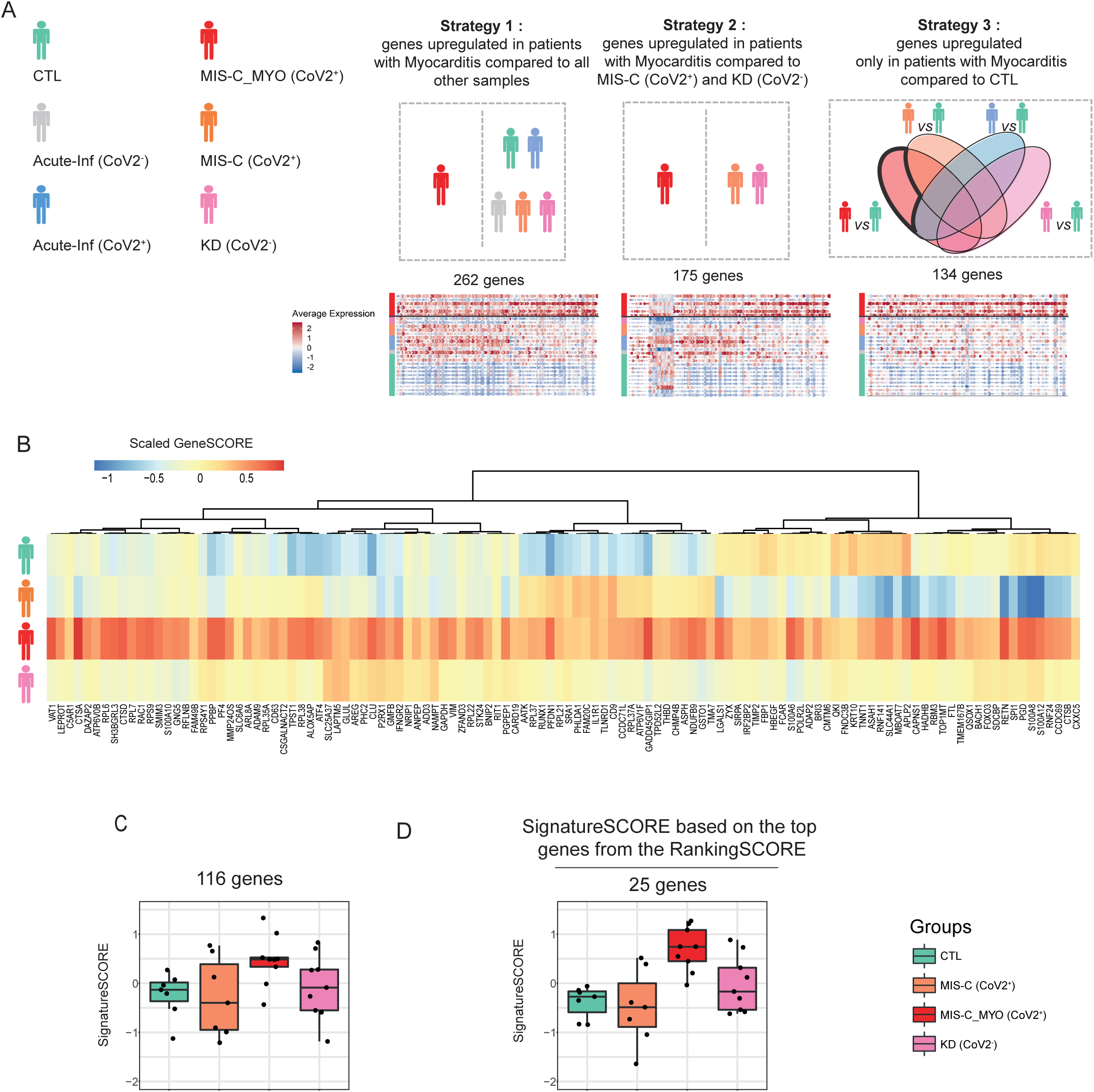
Identification of genes specifically upregulated in monocytes/dendritic cells of children of the MIS-C_MYO (CoV2^+^) group. **A.** Schematic representation of the different strategies used to extract 329 unique markers of the *MIS-C_MYO (CoV2^+^)* group from the monocytes/DCs clusters obtained from the single-cell dataset. Strategy 1: direct comparison of the monocytes/DCs cells of the *MIS-C_MYO (CoV2^+^)* group to all other samples. Strategy 2: direct comparison of the monocytes/DCs cells of *MIS-C_MYO (CoV2^+^)* to other samples with postacute hyperinflammation (*MIS-C (CoV2^+^)* and *KD (CoV2^-^))*. Strategy 3: selection of the genes upregulated only in the monocytes/DCs cells of *MIS-C_MYO (CoV2^+^)* when compared to CTL. Below each strategy, the corresponding dot plot obtained from SC-RNA-SEQ, with the number of upregulated genes. The average expression is represented by the centered scale expression of each gene. On the left, name of each group with its corresponding color is shown. **B.** Heatmap of expression of the 116/329 genes with a higher expression in *MIS-C_MYO (CoV2^+^)* than in other groups in the bulk dataset. Color scale indicates the scaled GeneSCORE (mean z-score of the gene in all samples of a group), with red and blue representing the highest and lowest expressions respectively. Hierarchical clustering of the genes was computed with a Pearson’s correlation as a distance. **C.** Box plot of the expression of the 116 genes validated in C, calculated as a SignatureScore. SignatureScore represents for each sample the mean z-score of the 116 genes selected in B in the bulk-RNA-SEQ dataset. (Explained in Figure S7A). **D.** Boxplot showing the SignatureScore computed on the expression of the top 25 genes, as ranked in Figure S7B, in the Bulk-RNA-SEQ dataset. **C & D.** Each dot represents a sample. Boxes range from the 25th to the 75th percentiles. The upper and lower whiskers extend from the box to the largest and smallest values respectively. Any samples with a value at most x1.5 the inter-quartile range of the hinge is considered an outlier and plotted individually.

## Discussion

Multi-parametric analysis of peripheral blood mononuclear cells from children with acute respiratory infection and postacute hyperinflammation, collected during the COVID-19 pandemic, allowed to detect an inflammatory profile associated with a loss of circulating monocytes and dendritic cells (DCs), as well as an upregulation of genes and pathways involving NF-κB signaling, oxidative stress with establishment of hypoxic conditions and VEGF signaling. These pathways were upregulated in both acute and postacute groups of patients, independently of SARS-CoV-2 infection. However, significant features of MIS-C with severe myocarditis were detected specifically in monocytes and DCs including low type-I and type-II IFN responses, decreased expression of NF-kB inhibitors, increased TNF-α signaling and overexpression of HIF-1α.

Acute cases were characterized by the detection of inflammatory markers in the plasma with a particularly strong elevation of IL-8 and CXCL1, two chemokines known to mediate neutrophil migration to the lung (Kunkel et al., 1991; Pease and Sabroe, 2002; Sawant et al., 2015) and a modest elevation of IFNα2 levels. These findings suggest that in some children, a suboptimal anti-viral type-I interferon response, alongside a hyperinflammatory response (IL-6 levels and exacerbation of the NF-kB pathway), could account for SARS-CoV-2 disease with pneumonia, as compared to the very usual benign or even asymptomatic clinical course of SARS-CoV-2 infection in children. This has been previously observed in severe Respiratory Syncytial Virus (RSV) infections (Hijano et al., 2019).

In the postacute patients, elevated levels of plasma IFN-γ, IFNα2, IL-10, IL-15, and, to a lesser extent, TNF-α, were found, as previously described in other cohorts (Brodsky et al., 2020; Carter et al., 2020; Consiglio et al., 2020; Esteve-Sole et al., 2021; Gruber et al., 2020). These findings are typical of an ongoing anti-viral immune response, not directly related to SARS-CoV-2 infection. In addition, elevated chemokines such as CCL2, CCL3 and CCL4 may recruit monocytes and DCs to tissues, possibly accounting for their reduced numbers observed in the blood of those patients. Additional mechanisms such as apoptosis or other cell death pathways may also be involved.

Cellular phenotypes that distinguish MIS-C from classical KD have been previously reported (Brodsky et al., 2020; Consiglio et al., 2020; Esteve-Sole et al., 2021). Brodin and colleagues described several key differences such as elevated IL-17, IL-6 and CXCL10 that were only observed in KD, associated with decreased naïve CD4^+^ T cells and increased central memory and effector memory CD4^+^ T cells in MIS-C. In the present study, high levels of IL-17, IL-6 and CXCL10, were both found in MIS-C and KD (CoV2^-^) groups. In addition, no major differences in CD4^+^ T cell compartments were detected. Accordingly, only a few differentially expressed genes were found between the MIS-C and KD (CoV2^-^) groups. These data support the hypothesis that MIS-C patients with KD features exhibit a molecular phenotype close to the one seen in KD patients, suggesting overlapping pathogenesis mechanisms (Gruber et al., 2020). Differences observed with previous reports by Brodin and colleagues, may be due to inclusion of only patients with criteria for complete or incomplete KD among the MIS-C cases, or technical differences in the respective studies, such as time of blood sampling relative to admission to hospital and medical treatments.

However, we did find noticeable differences in MIS-C cases with severe myocarditis with circulatory failure that required intensive care. The expression of a number of cytokines was further increased in these cases, most of them related to the NF-κB-TNF-α signaling axis. Elevated VEGF and TGF-α and TGF-β are potential drivers of angiogenesis and vascular homeostasis, whereas elevated chemokines (CCL2, CCL3, CCL20, CX3CL1, CXCL10) could mediate increased cell migration towards inflamed tissues. Molecular analysis confirmed an upregulation of genes belonging to the TNF-α and NF-κB signaling pathways that were specifically found in monocytes and DCs of MIS-C patients with severe myocarditis. A lower expression of NF-κB complex inhibitors, including *TNFAIP3* (A20), *TNFAIP2, NFKBIA, NFKBIZ,* was detected, suggesting a possible mechanism for NF-κB sustained activation which could then potentially lead to exacerbated TNF-α signaling. Overall, these results point to a potential role of monocytes and DCs in the pathogenesis of MIS-C with severe myocarditis, which might not be directly driven by SARS-CoV-2 infection, but rather due to a tolerance defect in limiting a pathological immune response, as already observed for other pathogens (Goodnow, 2021). It would be interesting to investigate the presence of genetic variants among MIS-C with severe myocarditis, in genes such as *TNFAIP3,* as already discussed (Goodnow, 2021). The apparent hypoxic conditions detected in children with myocarditis, could also account for the exacerbation of NF-κB signaling. HIF-1α, a sensor of oxidative stress, is well-known for being able to induce a switch from oxidative phosphorylation to glycolysis to limit generation of reactive oxygen species (ROS). It can also activate NF-κB signaling (D’Ignazio and Rocha, 2016; D’Ignazio et al., 2016). Additional environmental factors and/or genetic predispositions could also be involved. Another striking feature was the low expression of genes involved in type-I and type-II interferon responses, specifically in monocytes and DCs of children with myocarditis, although IFN-γ and IFNα2 proteins were elevated in the plasma of all MIS-C patients. This reduced response to type-I IFN in the most severe forms of MIS-C (with myocarditis and circulatory failure) is in part reminiscent of the impaired type-I IFN activity observed in the most severe forms of COVID-19 in adults (Bastard et al., 2020; Hadjadj et al., 2020; Zhang et al., 2020). The search for auto-antibodies against IFNα2 were negative (data not shown) but presence of autoantibodies to ISGs (interferon stimulated genes) cannot be excluded (Combes et al., 2021).

Overall, our findings depict a model, supported by previous publications (Amoah et al., 2015; Calabrese et al., 2004; Mann Douglas L., 2001), in which myocarditis is associated with an attenuated negative feedback loop of TNF-α-driven NF-κB activation, together with an excess of proangiogenic cytokines and chemokines that could attract activated myeloid and T cells to the myocardium tissue (**Figure S8**). Locally, it could lead to the production of inflammatory cytokines known to promote differentiation of cardiac fibroblasts into cardiac myofibroblasts (TNF-α, TGF-β, IL1-β, IL-13, IL-4, VEGF). Cardiac myofibroblasts, as previously reported, may secrete chemokines leading to further activation and recruitment of myeloid cells, creating a feed-forward loop of locally sustained inflammation and myocarditis (**Figure S8**) (Amoah et al., 2015; Angelo and Kurzrock, 2007; Delprat et al., 2020; Hua Xiumeng et al., 2020; Maloney and Gao, 2015).

Using SC-RNA-SEQ data, we defined a gene signature specific of SARS-CoV-2-related postacute hyperinflammatory illness with severe myocarditis that was further validated by a global transcriptomic analysis on PBMCs from an independent patient group. The genes defining this signature were consistently enriched in genes associated with inflammation, TNF-α and NF-κB signaling, oxidative stress and myocarditis (**Figure S7C**). Interestingly, among these genes, the S100 proteins and the calprotectin complex (S100A8/S100A9) in particular, were previously reported and proposed as biomarkers for the most severe adult form of COVID-19 with acute respiratory syndrome (**Figure S7D**) (Silvin et al., 2020).

Our study has several limitations, including the relatively low number of cases in each group, the lack of a comparison with asymptomatic or mildly symptomatic non-hospitalized children positive for SARS-CoV-2 and a longitudinal study of children with “classic” KD enrolled before the COVID-19 pandemic. All our cellular data were generated from frozen peripheral mononuclear cells, which does not allow a direct assessment of neutrophils. A parallel analysis of PMN (polymorphonuclear leukocytes) will be required. Endothelial and myocardiac cells are at least targets of the disease but may also contribute to the pathophysiology as described above. Also, additional data supporting gene expression findings will be necessary in future studies. Nevertheless, our study provides further in-depth molecular analysis of MIS-C with severe myocarditis. These severe forms were found to be associated with an excessive activation of the TNF-α, NF-κB signaling axis and poor response to type-I and type-II interferons in monocytes and DCs, secretion of cytokines promoting angiogenesis, chemotaxis and potential migration of activated myeloid cells and neutrophils in the myocardiac tissue. This may help to identify potential new clinical biomarkers and open new therapeutic strategies, including drugs targeting TNF-α or NF-κB pathways.

## Resource Availability

### Lead Contact

Further information and requests for resources and reagents should be directed to and will be fulfilled by the Lead Contacts, Mickaël Ménager and Frédéric Rieux-Laucat

### Material availability

This study did not generate new unique reagents.

### Data and Code availability

The SC-RNA-SEQ data have been deposited in the Gene Expression Omnibus (GEO) database under ID code GEO: GSE167029. The Bulk-RNA-SEQ data have been deposited under ID code GEO: GSE167028. Both can be accessed from the series code GEO: GSE167030

## Experimental Model and Subject details

### Patients and definitions

This prospective multicenter cohort study included children (age ≤ 18 years at the time of admission) suspected of infection with SARS-CoV-2 between April 6, 2020 and May 30, 2020. Clinical aspects of 22 of the included patients were previously reported (Toubiana et al., 2020, 2021). Children admitted with fever in general pediatric wards or pediatric intensive care units of Tertiary French hospitals involved in the research program, suspected of SARS-CoV-2 related illness and who underwent routine nasopharyngeal swabs for SARS-CoV-2 RT-PCR (R-GENE, Argene, Biomerieux, Marcy l’Etoile) or SARS-CoV-2 IgG serology testing (Architect SARS-CoV-2 chemiluminescent microparticle immunoassay; Abbott Core Laboratory, IL, USA), were eligible. The study was approved by the Ethics Committee (Comité de Protection des Personnes Ouest IV, n° DC-2017-2987). All parents provided written informed consent.

Case definition for pediatric COVID-19 acute infection was presence of fever, fatigue, neurological abnormalities, gastro-intestinal or respiratory signs, associated with a concomitant nasopharyngeal swab positive for SARS-CoV-2 RT-PCR, and absence of MIS-C criteria (Zimmermann and Curtis, 2020). Case definition for postacute hyperinflammatory illness (**Figure 1**) was presence of fever, laboratory evidence of inflammation and clinically severe illness with multisystem involvement, during the SARS-CoV-2 epidemic period (Datta et al., 2020). This may include children with features of KD; criteria of the American Heart Association was used to define for complete (Fever > 4 days and ≥ 4 principal criteria) or incomplete KD (Fever > 4 days and 2 or 3 principal criteria, and without characteristics suggestive of another diagnosis) (McCrindle et al., 2017). Among cases with postacute hyperinflammatory illness, children with a positive SARS-CoV-2 testing (RT-PCR or serology) were considered to have MIS-C according to CDC and WHO criteria to define MIS-C (CDC, 2020). Patients with postacute hyperinflammatory illness, negative SARS-CoV-2 testing (RT-PCR or serology), and criteria for KD, were considered as patients with KD-like illness. Patients with MIS-C with clinical signs of circulatory failure requiring intensive care, with elevated high-sensitivity cardiac troponin I levels (>26 ng/mL) and /or decreased cardiac function (diastolic or systolic ventricular dysfunction at echocardiography), were considered to have MIS-C with severe myocarditis (Brissaud et al., 2016; Canter Charles E. and Simpson Kathleen E., 2014).

For each included patient, we collected demographic data, symptoms, results of SARS-CoV-2 testing and other laboratory tests, echocardiograms, and treatments. All patients with negative initial serology testing were retested after an interval of at least 3 weeks (Architect SARS-CoV-2 chemiluminescent microparticle immunoassay; Abbott Core Laboratory).

Healthy controls were recruited before the COVID-19 pandemic (before November 2019).

### Samples

For each patient and healthy donor, peripheral blood samples were collected on EDTA and lithium heparin tubes. After a centrifugation of the EDTA tube at 2300rpm for 10 minutes, plasma was taken and stored at -80°C before cytokine quantification. PBMCs were isolated from the lithium heparin samples, frozen as described below and stored at -80°C and were used for both bulk and single-cell RNAseq, as well as cell phenotyping by CyTOF. The workflow is summarized in Figure 1B.

### Isolation of PBMCs

Peripheral blood samples were collected on lithium heparin. PBMCs were isolated by density gradient centrifugation (2200 rpm without break for 30 minutes) using Ficoll (Eurobio Scientific, Les Ulis, France). After centrifugation, cells were washed with Phosphate-buffered saline (PBS) (Thermo Fisher scientific, Illkirch, France). The pellet was resuspended in PBS and cells were centrifuged at 1900 rpm for 5 minutes. Finally, the PBMCs pellet was frozen in a medium containing 90% of Fetal Bovine Serum (FBS) (Gibco, Thermo Fisher scientific, Illkirch, France) and 10% of dimethyl sulfoxide (DMSO) (Sigma Aldrich, St. Quentin Fallavier, France).

### Cytokine measurements

Prior to protein analysis plasma samples were treated in a BSL3 laboratory for viral decontamination using a protocol previously described for SARS-CoV (Darnell and Taylor, 2006), which we validated for SARS-CoV-2. Briefly, samples were treated with TRITON X100 (TX100) 1% (v/v) for 2hrs at Room Temperature. IFNα2, IFNγ, IL-17A, (triplex) and IFNβ (single plex) protein plasma concentrations were quantified by Simoa assays developed with Quanterix Homebrew kits as previously described (Rodero et al., 2017). The limit of detection of these assays were 0.6 pg/mL for IFNβ, 2 fg/mL for IFNα2, 0.05 pg/ml for IFNγ and 3 pg/mL for IL17A including the dilution factor. IL-6, TNFα, and IL-10 were measured with a commercial triplex assay (Quanterix). Additional plasma cytokines and chemokines (44 analytes) were measured with a commercial Luminex multi-analyte assay (Biotechne, R&D systems).

### Serology assays

SARS-CoV-2 specific antibodies were quantified using assays previously described (Grzelak et al., 2020). Briefly, a standard ELISA assay using as target antigens the extracellular domain of the S protein in the form of a trimer (ELISA tri-S) and the S-Flow assay, which is based on the recognition of SARS-CoV-2 S protein expressed on the surface of 293T cells (293T-S), were used to quantify SARS-CoV-2 specific IgG and IgA subtypes in plasma. Assay characteristics including sensitivity and specificity were previously described (Grzelak et al., 2020).

### Cell Phenotyping

To perform high-dimensional immune profiling of PBMCs, we used the Maxpar^®^ Direct™ Immune Profiling System (Fluidigm, Inc France) with a 30-marker antibody panel, for CyTOF (Cytometry by Time Of Flight). Briefly, 3×10^6^ PBMCs resuspended in 300 µl of MaxPar Cell Staining Buffer were incubated for 20 minutes at room temperature after addition of 3 μL of 10 KU/mL heparin solution and 5 µl of Human TruStain FcX (Biolegend Europ, Netherland). Then 270 μL of the samples were directly added to the dry antibody cocktail for 30 minutes. 3 mL of MaxPar Water was added to each tube for an additional 10-min incubation. Three washes were performed on all the samples using MaxPar Cell Staining Buffer and they were fixed using 1.6% paraformaldehyde (Sigma-Aldrich, France). After one wash with MaxPar Cell Staining Buffer, cells were incubated one hour in Fix and Perm Buffer with 1:1000 of Iridium intercalator (pentamethylcyclopentadienyl-Ir (III)-dipyridophenazine, Fluidigm, Inc France). Cells were washed and resuspended at a concentration of 1 million cells per mL in Maxpar Cell Acquisition Solution, a high-ionic-strength solution, and mixed with 10% of EQ Beads immediately before acquisition.

Acquisition of the events was made on the Helios mass cytometer and CyTOF software version 6.7.1014 (Fluidigm, Inc Canada) at the “Plateforme de Cytométrie de la Pitié-Salpetriere (CyPS).” An average of 500,000 events were acquired per sample. Dual count calibration, noise reduction, cell length threshold between 10 and 150 pushes, and a lower convolution threshold equal to 10 were applied during acquisition. Mass cytometry standard files produced by the HELIOS were normalized using the CyTOF Software v. 6.7.1014. For data cleaning, 4 parameters (centre, offset, residual and width) are used to resolve ion fusion events (doublets) from single events from the Gaussian distribution generated by each event (Bagwell et al., 2020). Subsequent to data cleaning, the program produces new FCS files consisting of only intact live singlet cells. These data were analyzed in FlowJo v10.7.1 using 3 plugins (DownSampleV3, UMAP and FlowSOM) with R v4.0.2. To increase efficiency of the analysis, samples were downsampled to 50 000 cells, using the DownSample V3 plugin. All samples were concatenated and analyzed in an unsupervised manner. Anti-CD127 antibody had to be excluded due to poor staining. Clustering was performed using FlowSOM (Van Gassen et al., 2015). The number of clusters was set to forty-five in order to overestimate the populations and detect smaller subpopulations. Grid size of the self-organizing map was set to 20×20. Resulting clusters were annotated as cell populations following the kit manufacturer’s instruction. When several clusters were identified as the same cell types, they were concatenated into a single cell population. For visualization purposes, UMAP was computed with the UMAP pluggin (McInnes et al.) with the following parameters: metric (Euclidean), nearest neighbors (15), minimum distance (0.5) and number of components (2).

### Single-cell transcriptomic (SC-RNA-SEQ)

SC-RNA-SEQ analyses were performed on frozen PBMCs isolated from heparin blood samples. PBMCs were thawed according to 10X Genomics protocol. The SC-RNA-SEQ libraries were generated using Chromium Single Cell 3′ Library & Gel Bead Kit v.3 (10x Genomics) according to the manufacturer’s protocol. Briefly, cells were counted, diluted at 1000 cells/µL in PBS+0,04% and 20,000 cells were loaded in the 10x Chromium Controller to generate single-cell gel-beads in emulsion. After reverse transcription, gel-beads in emulsion were disrupted. Barcoded complementary DNA was isolated and amplified by PCR. Following fragmentation, end repair and A-tailing, sample indexes were added during index PCR. The purified libraries were sequenced on a Novaseq 6000 (Illumina) with 28 cycles of read 1, 8 cycles of i7 index and 91 cycles of read 2.

Sequencing reads were demultiplexed and aligned to the human reference genome (GRCh38, release 98, built from Ensembl sources), using the CellRanger Pipeline v3.1. Unfiltered RNA UMI counts were loaded into Seurat v3.1 (Stuart et al., 2019) for quality control, data integration and downstream analyses. Apoptotic cells and empty sequencing capsules were excluded by filtering out cells with fewer than 500 features or a mitochondrial content higher than 20%. Data from each sample were log-normalized and scaled, before batch correction using Seurat’s FindIntegratedAnchors. For computational efficiency, anchors for integration were determined using all control samples as reference and patient samples were projected onto the integrated controls space. On this integrated dataset, we computed the principal component analysis on the 2000 most variable genes. UMAP was carried out using the 20 most significant principal components (PCs), and community detection was performed using the graph-based modularity-optimization Louvain algorithm from Seurat’s FindClusters function with a 0.8 resolution. Cell types labels were assigned to resulting clusters based on a manually curated list of marker genes as well as previously defined signatures of the well-known PBMCs subtypes (Monaco et al., 2019). Despite filtering for high quality cells, five clusters out of the twenty-six stood out as poor quality clusters and were removed from further analysis, namely: one erythroid-cell contamination; one low UMI cluster from a single control; two clusters of proliferating cells originating from a patient with EBV co-infection and one megakaryocytes cluster. In total 152,201 cells were kept for further analysis.

After extraction and reclustering of high-quality cells, differential expression was performed separately on all PBMCs, Monocytes/DCs, T cells or B cells. Differential expression testing was conducted using the FindMarkers function of Seurat on the RNA assay with default parameters. Genes with log(FC) > 0.25 and adjusted p-values ≤0.05 were selected as significant.

### Bulk RNA-sequencing (Bulk-RNA-SEQ)

Bulk-RNA-SEQ analyses were performed on frozen PBMCs extracted from heparin samples. RNA was extracted from PBMCs following the instructions of RNeasyR Mini kit (Qiagen, Courtaboeuf, France). To note, the optional step with the DNase was performed. RNA integrity and concentration were assessed by capillary electrophoresis using Fragment Analyzer (Agilent Technologies). RNAseq libraries were prepared starting from 100 ng of total RNA using the Universal Plus mRNA-Seq kit (Nugen) as recommended by the manufacturer. The oriented cDNA produced from the poly-A+ fraction was sequenced on a NovaSeq6000 from Illumina (Paired-End reads 100 bases + 100 bases). A total of ∼50 millions of passing-filters paired-end reads was produced per library.

Paired-end RNA-seq reads were aligned to the human Ensembl genome GRCh38.91 using Hisat2 (v2.0.4)(Kim et al., 2019) and counted using featureCounts from the Subread R package. The raw count matrix was analyzed using DESeq2 (version 1.28.1) (Love et al., 2014). No pre-filtering was applied to the data. Differential expression analysis was performed using the “DESeq” function with default parameters. For visualization and clustering, the data was normalized using the ‘variant stabilizing transformation’ method implemented in the “vst” function. Plots were generated using ggplot2 (version 3.3.2), and pheatmap (version 1.0.12). During exploratory analyses, it was noted that the clustering was mainly driven by the sex of the patients. To remove this effect, it was included in the regression formula for DESeq (∼sex + groups), and then removed following vst transformation, using “removeBatchEffect” from the “limma” package (version 3.44.3).

### Gene signature analysis

To identify genes that could be used as markers of severe myocarditis in the SC-RNA-SEQ dataset, three initial strategies were used, all based on differential expression and selection of the upregulated genes. First, we performed the differential expression between *MIS-C_MYO (CoV2^+^)* samples and all other samples. Second, differential analysis was computed between *MIS-C_MYO (CoV2^+^)* and other samples with postacute hyperinflammatory illness. In the last strategy, we selected genes that were upregulated between the *MIS-C_MYO (CoV2^+^)* and the CTL, but not upregulated in any other group compared to the CTL (Figure 7A). These three strategies allowed us to identify 329 unique genes.

To further explore whether these genes could be considered as markers of severe myocarditis, we analyzed their expression profile in our bulk RNA-SEQ dataset. This dataset excluded samples from patients of the *MIS-C_MYO (CoV2^+^)* that were included in the SC-RNA-SEQ cohort. Vst-transformed counts were log2-normalized and converted to z-score using the scale function in R (v 4.0.2). A GeneSCORE was computed for each group as the mean z-score of the samples of a group. Heatmaps representing this GeneSCOREgroup were performed using pheatmap. Hierarchical clustering of the 329 previously identified genes was performed using the complete method on the distance measured using Pearson’s correlation, as implemented by pheatmap. The hierarchical clustering was divided into 15 main clusters, 4 of which had the expected pattern of expression: Clusters that had a higher expression in *MIS-C_MYO (CoV2^+^)* than any other group were selected, resulting in 116 genes. A signature score for each sample was performed on these genes, corresponding to the mean expression (z-score) of these N genes in each sample (SignatureSCORE).

These genes were subsequently ranked based on the following equation:

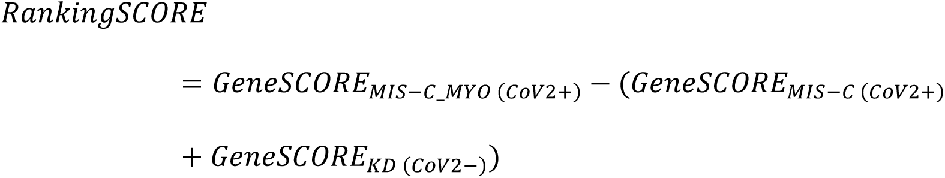

where the SCOREs represent the mean expression (z-score) in each disease groups, and the SignatureScore was computed on the top 25 genes.

### Quantification and statistical analysis

Cytokine heatmaps were made with Qlucore OMICS explore (version 3.5(26)) and dot plots with GraphPad Prism (version 8). Differentially secreted cytokines were included in the heat maps based on a 1.5 Fold Change (FC) comparison between groups as indicated. Dot plot differences between each group were identified by Kruskal-Wallis tests followed by post-hoc multiple comparison Dunn’s test.

Statistical tests for cellular composition analysis in both the CyTOF and SC-RNA-SEQ datasets were performed in R v3.6.1. Kruskal-Wallis test followed by post-hoc multiple comparison Dunn’s test was applied to assess differences in cell population proportions (*: p ≤0.05; **: p ≤0.01; ***: p ≤0.001).

Differential expression testing in the SC-RNA-SEQ dataset was conducted using the FindMarkers function in Seurat, with default Wilcoxon testing. P-values were controlled using Bonferroni correction. Genes with an absolute log(fold-change) ≥0.25 and an adjusted p-value ≤0.05 were selected as differentially expressed. Pathways analysis was performed using both the Ingenuity pathway analysis v57662101 software (IPA (QIAGEN Inc.) and EnrichR (Chen et al., 2013; Kuleshov et al., 2016). Heatmaps were extracted from the comparison module in IPA. Pathways with an absolute z-score lower than 2 or a Bonferroni-Hochberg corrected p-values higher than 0.05 were filtered out. Reactome 2016 and Molecular Signature DataBase Hallmark 2020 (MSigDB Hallmark 2020) pathway enrichment analysis were performed using EnrichR. The TRRUST transcription factors 2019 (Han et al., 2018) used for the transcription factors enrichment analysis was performed using Enrich R.

## Author contributions

CdC, ML, SM, AM, NS, FC, VGP, LB generated and analyzed data. MB, AB, BPP, GA, TF analyzed data. MB, LG, PG, JDS, HM, OS, CB, PB, and JLC generated data. CdC, ML, AF design figures and wrote manuscript. DD, FRL, JT and MMM conceived the study, analyzed data, wrote the manuscript and supervised the study.

## Acknowledgements

The study was supported by the Institut National de la Santé et de la Recherche Médicale (INSERM), by the “URGENCE COVID-19” fundraising campaign of Institut Pasteur, by the Atip-Avenir, Emergence ville de Paris program and fond de dotation Janssen Horizon and by government grants managed by the Agence National de la Recherche as part of the “Investment for the Future” program (Institut Hospitalo-Universitaire Imagine, grant ANR-10-IAHU-01, Recherche Hospitalo-Universitaire, grant ANR-18-RHUS-0010, Laboratoire d’Excellence ‘‘Milieu Intérieur”, grant ANR-10-LABX-69-01), the Centre de Référence Déficits Immunitaires Héréditaires (CEREDIH), the Agence National de la Recherche (ANR-flash Covid19 “AIROCovid” to FRL and “CoVarImm” to DD and JDS), and by the FAST Foundation (French Friends of Sheba Tel Hashomer Hospital). The LabTech Single-Cell@Imagine is supported by the Paris Region and the “Investissements d’avenir” program through the 2019 ATF funding – Sésame Filières PIA (Grant N°3877871).

CdC is the recipient of a CIFRE-PhD (Sanofi). L.B. was a recipient of an Imagine institute PhD international program supported by the Fondation Bettencourt Schueller. L.B. was also supported by the EUR G.E.N.E. (reference #ANR-17-EURE-0013) and is part of the Université de Paris IdEx #ANR-18-IDEX-0001 funded by the French Government through its “Investments for the Future” program. S.M. was a recipient of an INSERM and Institut Imagine post-doctorat program supported by the Fondation pour la Recherche Médicale (FRM N°SPF20170938825). NS was a recipient of the Pasteur-Roux-Cantarini Fellowship. VGP obtained an Imagine international PhD fellowship program supported by the Fondation Bettencourt Schueller. BPP is the recipient of an ANRS post-doctoral fellowship. We thank *Imagine* genomic, bioinformatic and single-cell core facilities, the Institut Pasteur Cytometry and Biomarkers UTechS platform and the Pitié-Salpêtrière Cytometry platform CyPS.

---

**Table S1.**
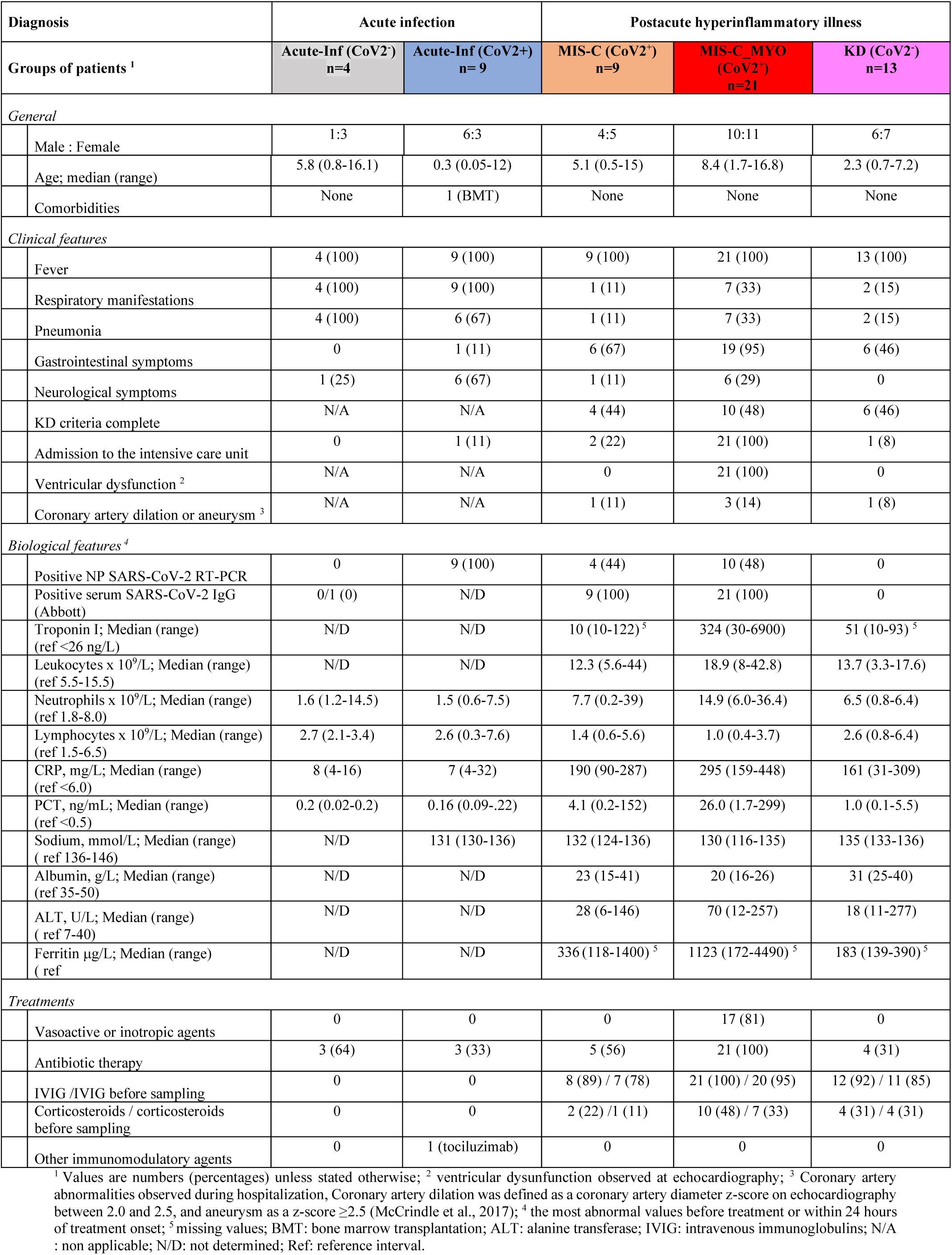
Clinical and biological features of pediatric patients enrolled. All children and adolescents included in the study were suspected of SARS-CoV-2 illness between April 6, 2020 and May 30, 2020 and displayed either an acute respiratory infection or a postacute inflammatory illness.

**Figure S1:**
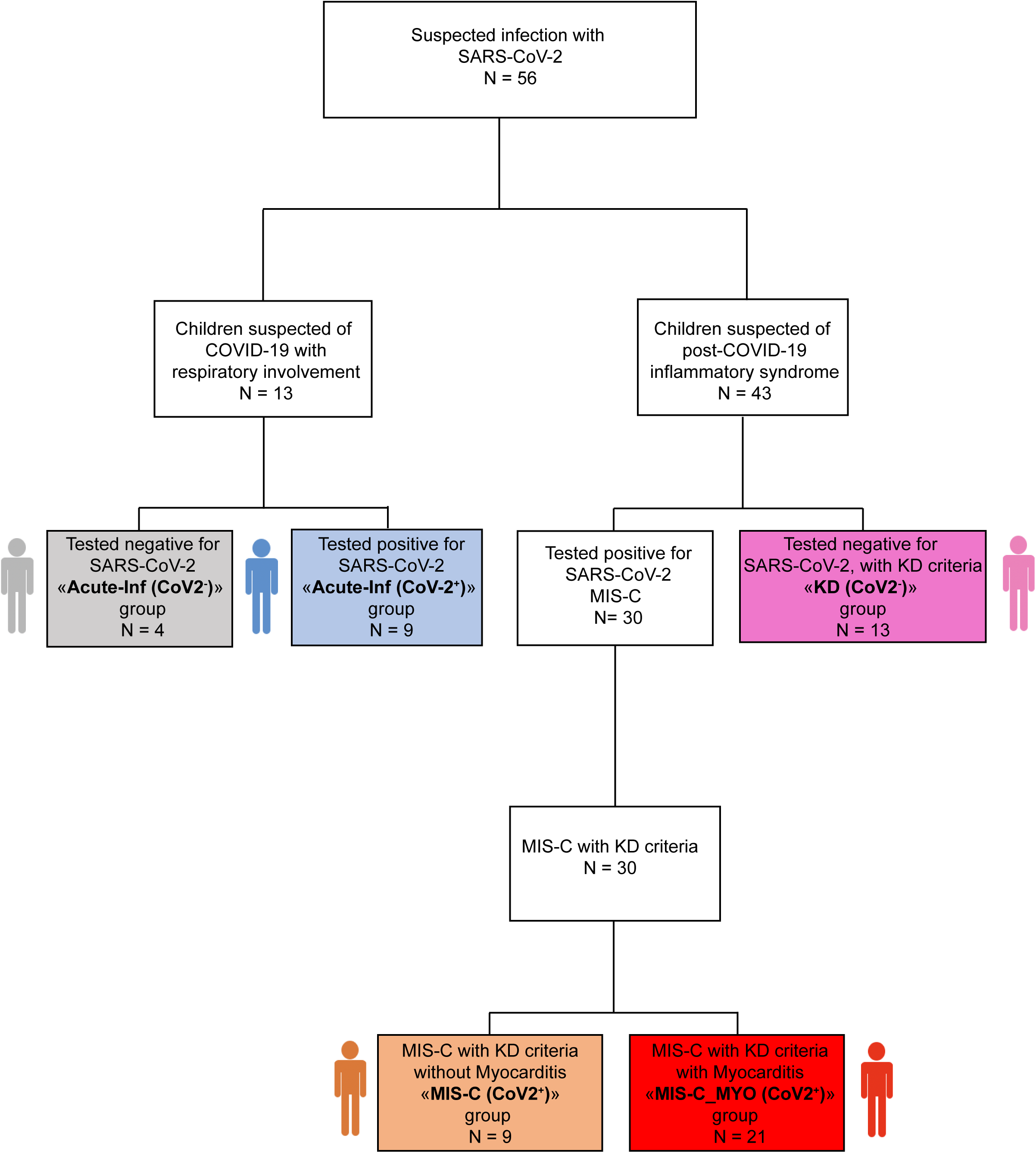
Flowchart describing the patients enrolled. Related to Figure 1. Each group with a name in bold and a color associated corresponds to a group of patients analyzed in this study. Names and colors are used throughout the manuscript. “N” represents the number of patients enrolled in each group.

**Figure S2:**
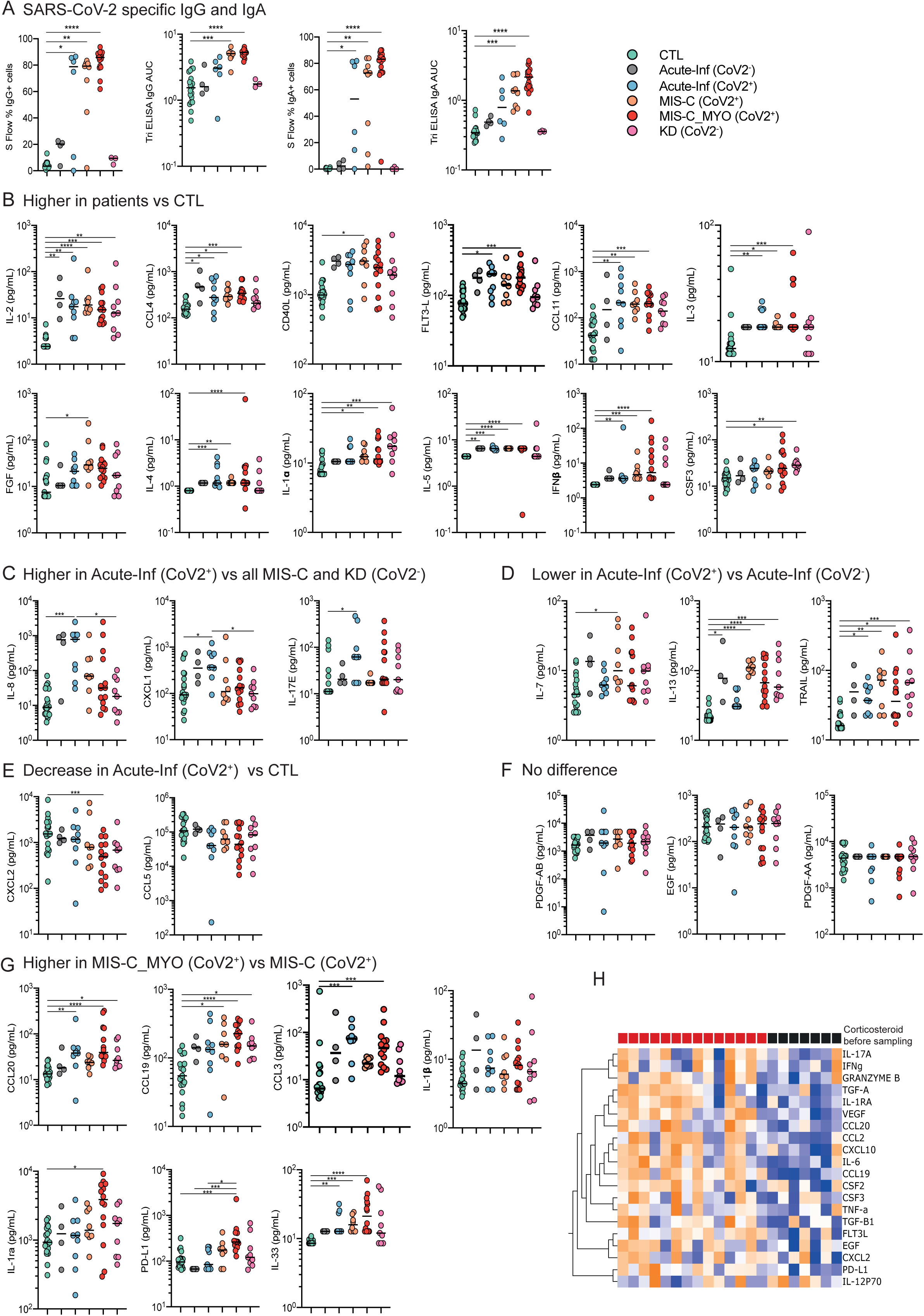
Immunoglobulin and cytokine analyses. Related to Figure 2. **A.** SARS-CoV-2 specific immunoglobulins (IgG, left panels; IgA, right panels) were dosed in the plasma of the different groups, by two methods: S Flow and Tri ELISA. **B - G.** Dot plots of cytokines differentially expressed and grouped based on the description of each sub panel. ρ values are calculated by Kruskall-Wallis test for multiple comparisons, followed by a post hoc Dunn’s test. *(ρ < 0.05), **(ρ < 0.01), ***(ρ < 0.001). **H.** Hierarchical clustering of cytokines differentially expressed among the patients of the *MIS-C_MYO (CoV2^+^)* group without (red square) or with glucocorticosteroid treatment (black square) before blood sampling.

**Figure S3:**
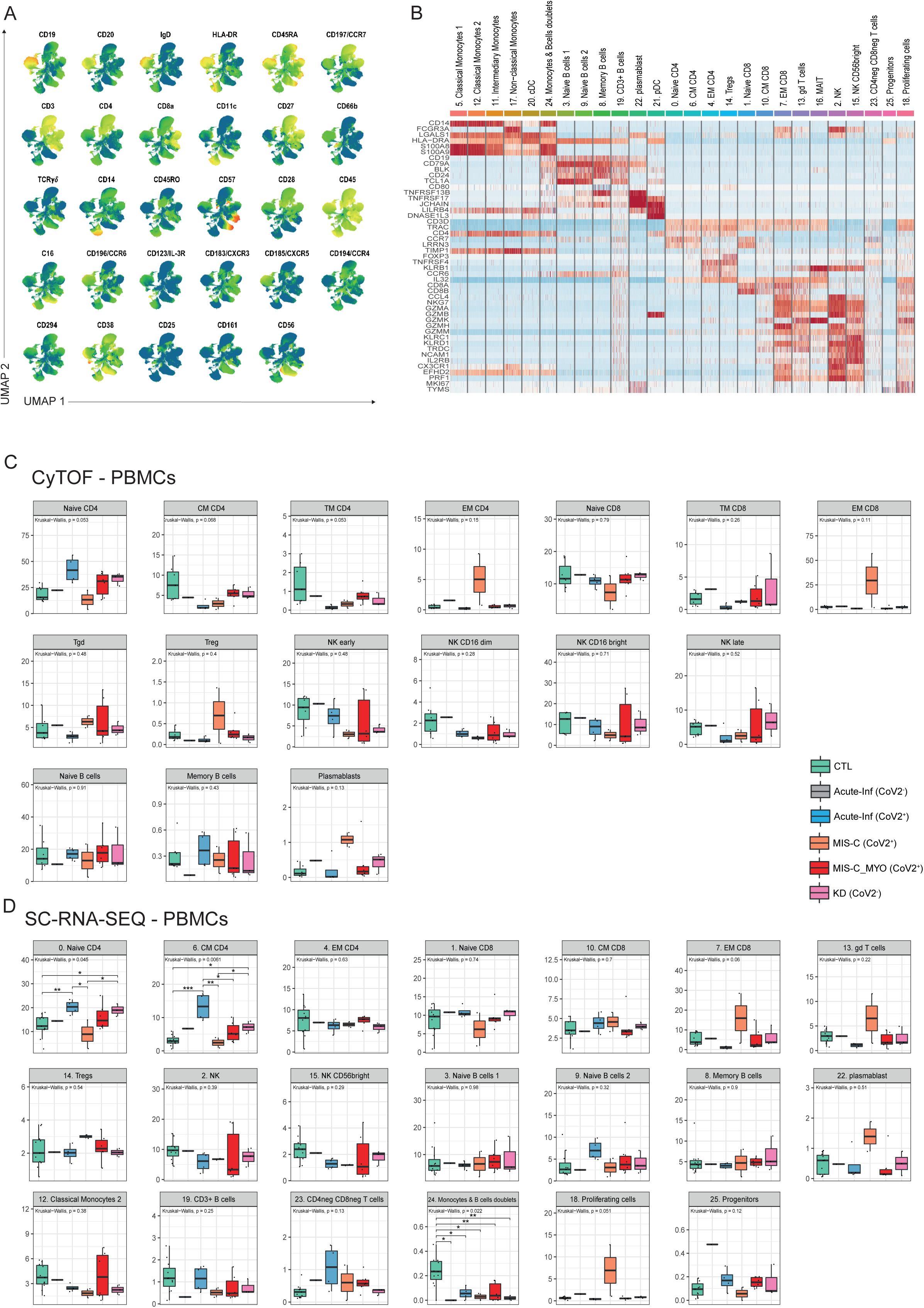
Identification of cell clusters and PBMCs distribution. Related to Figure 3. **A.** UMAP showing all cell surface proteins measured by CyTOF to identify clusters displayed in Figure 3A. **B.** Heatmap of the genes (y axis) used in SC-RNA-SEQ to identify clusters of the UMAP in Figure 3B. Color scale represents scaled expression of the genes. Red and blue indicate high and low expression respectively. **C & D.** Boxplots showing the percentage of cell populations from CyTOF (C) and SC-RNA-SEQ (D). Among the *MIS-C_(CoV2^+^)* group, one patient was diagnosed with a primary Epstein Barr viral infection, characterized at the cellular level by an increase of CD4, CD8 effector memory T cells, Tregs and also proliferating cells. (CTL, green; *Acute-inf (CoV2^-^)*, gray; *Acute-inf (CoV2^+^)*, blue; *MIS-C (CoV2^+^)*, orange; *MIS-C_MYO (CoV2^+^)*, red; *KD (CoV2^-^)*, pink). In the boxplots, each dot represents a sample. Boxes range from the 25th to the 75th percentiles. The upper and lower whiskers extend from the box to the largest and smallest values respectively. Any samples with a value at most x1.5 the inter-quartile range of the hinge is considered an outlier and plotted individually. ρ values are calculated by Kruskal-Wallis test for multiple comparison, followed by a post hoc Dunn’s test. *(ρ < 0.05), **(ρ < 0.01), ***(ρ < 0.001).

**Figure S4:**
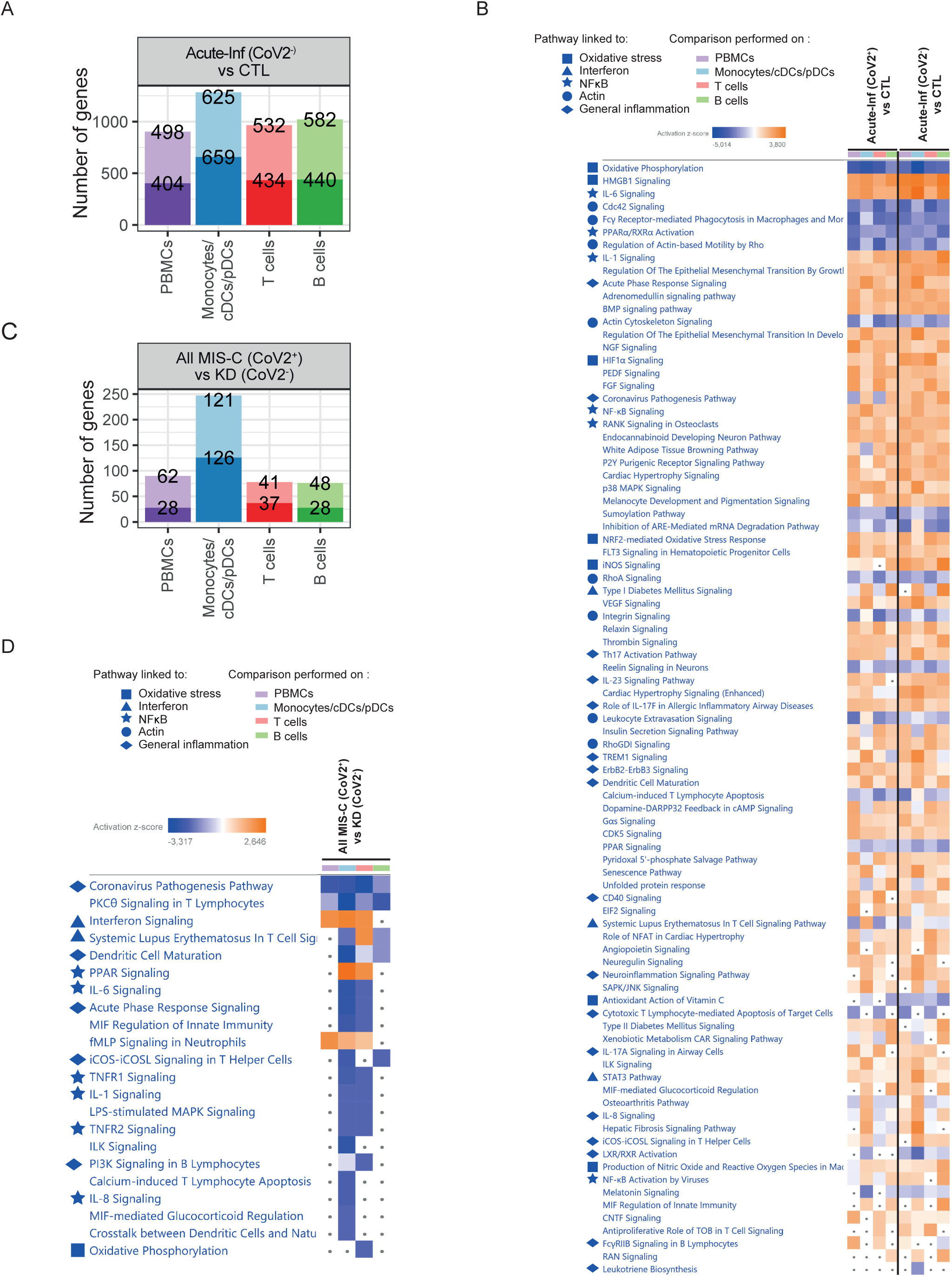
Genes and pathways differentially regulated from SC-RNA-SEQ in acute infection and postacute hyperinflammation when comparing patients with or without evidence of SARS-CoV-2 infection. Related to Figure 4. **A.** Bar charts of the number of up- and downregulated genes in *Acute-inf (CoV2^-^)* compared to CTL, in PBMCs, monocytes/DCs, T and B cell clusters obtained following SC-RNA-SEQ experiments as displayed in Figure 3B. Heatmap of the canonical pathways obtained with IPA from DEGs of *Acute-inf (CoV2^+^)*, as shown in 4A or of *Acute-inf (CoV2^-^)*, S4A, compared to CTL in PBMCs, monocytes/DCs, T and B cells. **C.** Bar charts of the number of up- and downregulated genes in *All MIS-C (CoV2^+^)* compared directly to *KD (CoV2^-^)*, in PBMCs, monocytes/DCs, T and B cells. **D.** Heatmap of the canonical pathways obtained with IPA from DEGs of *All MIS-C (CoV2^+^)* compared directly to *KD (CoV2^-^)* in PBMCs, monocytes/DCs, T and B cells. **A & C.** The top value on the light-colored bars represent the upregulated genes and the bottom dark represent the downregulated genes. **B & D.** Symbols are used in front of the pathways to represent pathways belonging to the same functional groups. Pathways with an absolute z-score ≤ 2 or adjusted p-value > 0.05 in all conditions were filtered out. Z-score > 2 means that a function is significantly increased (orange) whereas a Z-score < −2 indicates a significantly decreased function (blue). Grey dots indicate non-significant pathways (ρ > 0.05).

**Figure S5:**
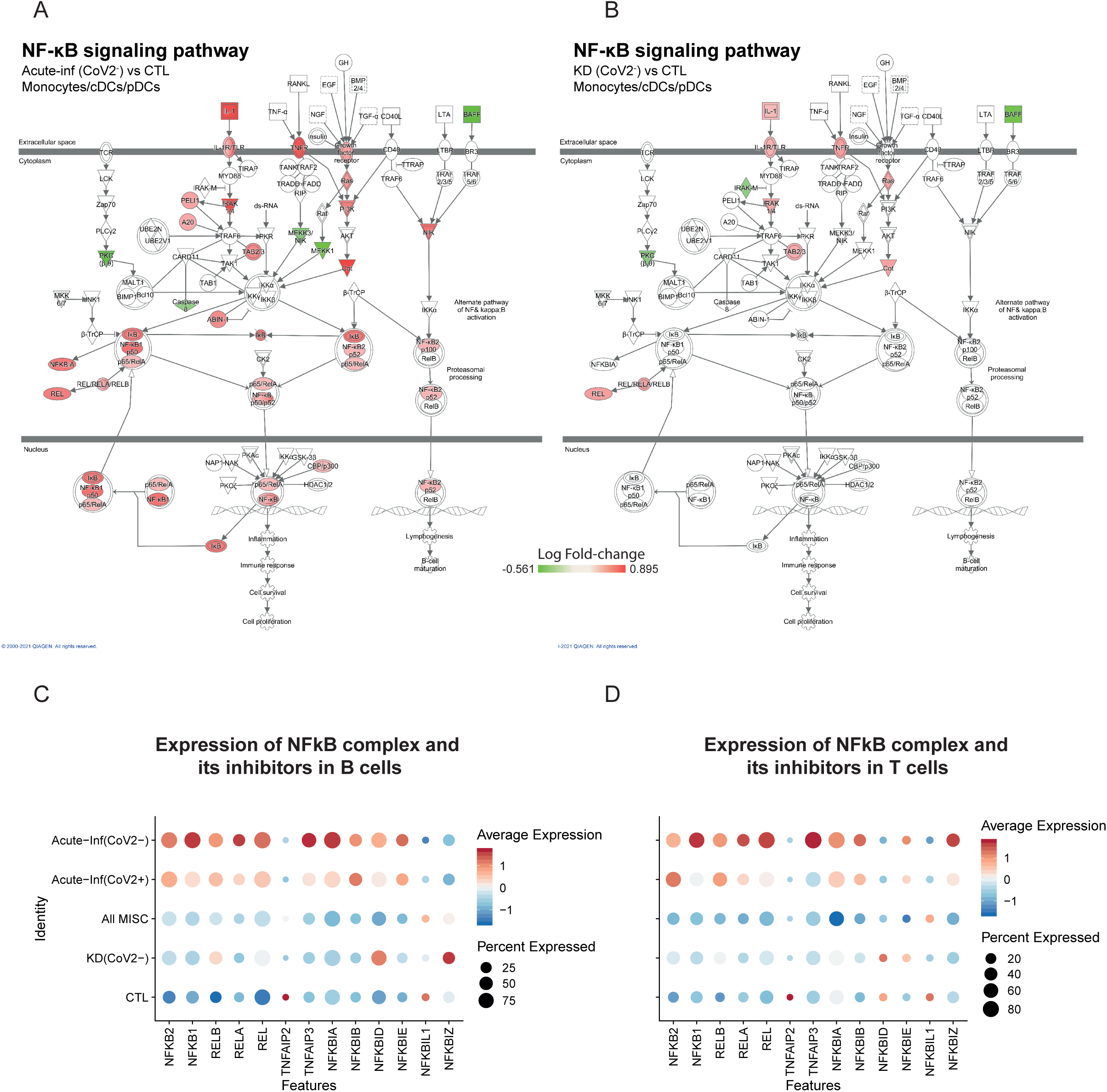
NF-κB signaling pathway activation in acute infection and postacute hyperinflammatory illness unrelated to SARS-CoV2 infection. Related to Figure 5. **A & B.** NF-κB signaling pathway from the IPA software, displayed in the monocytes/DCs cells in *Acute-inf (CoV2^-^)* versus CTL (A) or *KD (CoV2^-^)* versus CTL (B). Changes in gene expression are represented by the log fold change (increase in red and decrease in green). **C & D.** Dot plot of the expression in B cells (C) or T cells (T) of the positive (from *NFKB2 to REL*) and negative regulators (from *TNFAIP2* to *NFKBIZ*) of NF-κB complex in all disease groups.

**Figure S6:**
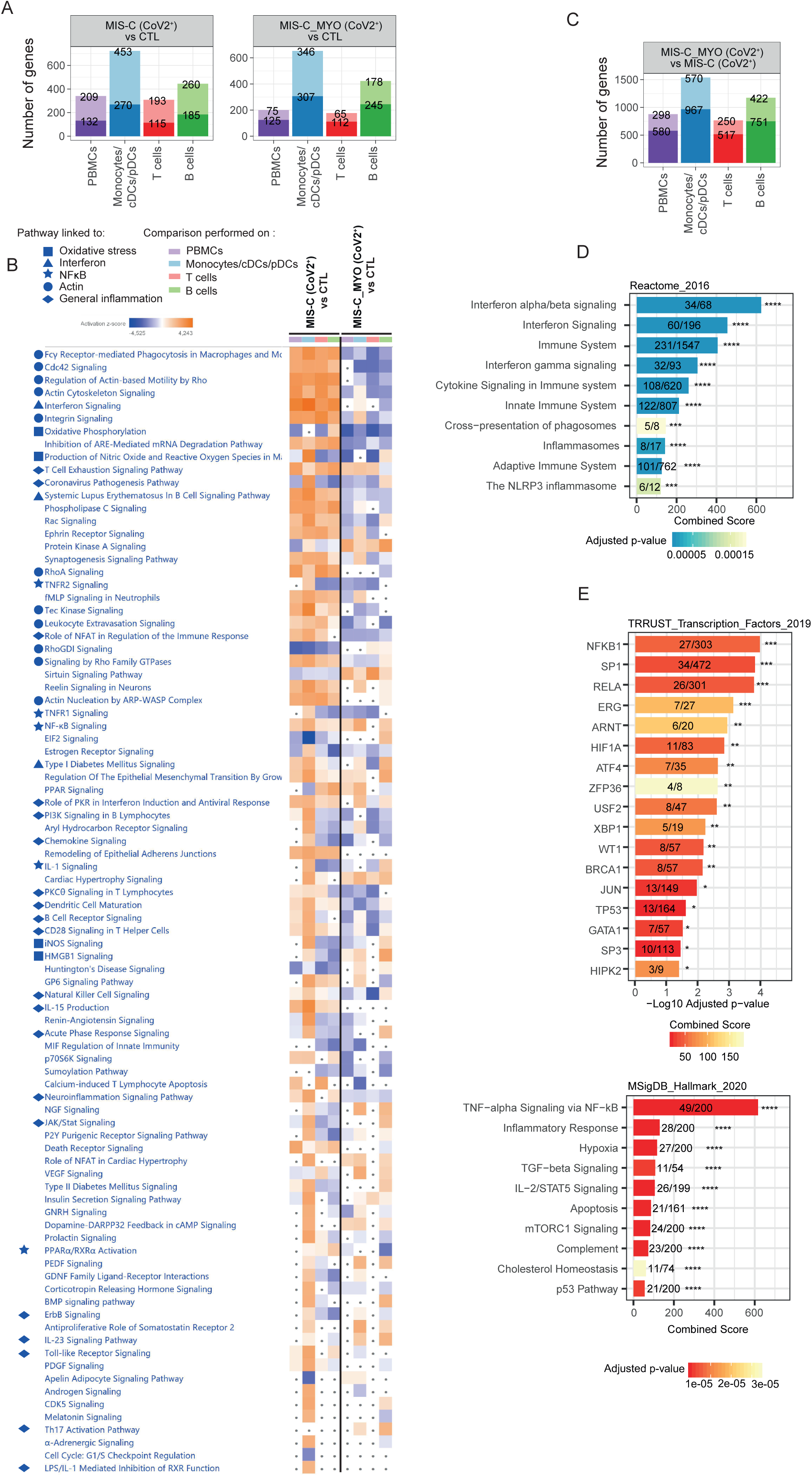
Pathway enrichment analysis in MIS-C (CoV2^+^) with or without severe myocarditis. Related to Figure 6. **A.** Bar charts of the number of up- and downregulated genes in *MIS-C (CoV2^+^)* (left) and *MIS-C_MYO (CoV2^+^)* (right) compared to CTL in PBMCs, monocytes/DCs, T and B cells. **B.** Heatmap of the canonical pathways obtained with IPA from DEGs of *MIS-C (CoV2^+^)* or *MIS-C_MYO (CoV2^+^)* compared to CTL in PBMCs, monocytes/DCs, T and B cells. Symbols are used in front of the pathways to represent pathways belonging to the same functional groups. Pathways with an absolute z-score ≤ 2 or adjusted p-value > 0.05 in all conditions were filtered out. Z-score > 2 means that a function is significantly increased (orange) whereas a Z-score < −2 indicates a significantly decreased function (blue). Grey dots indicate non-significant pathways (ρ > 0.05). **C.** Bar charts of the number of up- and downregulated genes in *MIS-C_MYO (CoV2^+^)* compared to *MIS-C (CoV2^+^)* in PBMCs, monocytes/DCs, T and B cells. **D.** Bar charts of the top 10 pathways from Reactome_2016, predicted in EnrichR to be modulated by the downregulated genes in the monocytes/DCs cells of *MIS-C_MYO (CoV2^+^)* compared to *MIS-C (CoV2^+^)*. Pathways are ranked based on combined score. **E.** Bar charts of the Transcription factors from TRRUST database (top) and the top 10 pathways from MSigDB_Hallmark_2020 (bottom) predicted in EnrichR to be modulated by the genes upregulated in monocytes/DC of *MIS-C_MYO (CoV2^+^)* compared to *MIS-C (CoV2^+^)*. Pathways are ranked based on combined score; Transcription factors are ranked based on adjusted p-values. **A & C.** The top values on the light-colored bars represent the upregulated genes and the bottom dark represent the downregulated genes. **D & E.** *(ρ < 0.05), **(ρ < 0.01), ***(ρ < 0.001), ****(ρ < 0.0001)

**Figure S7:**
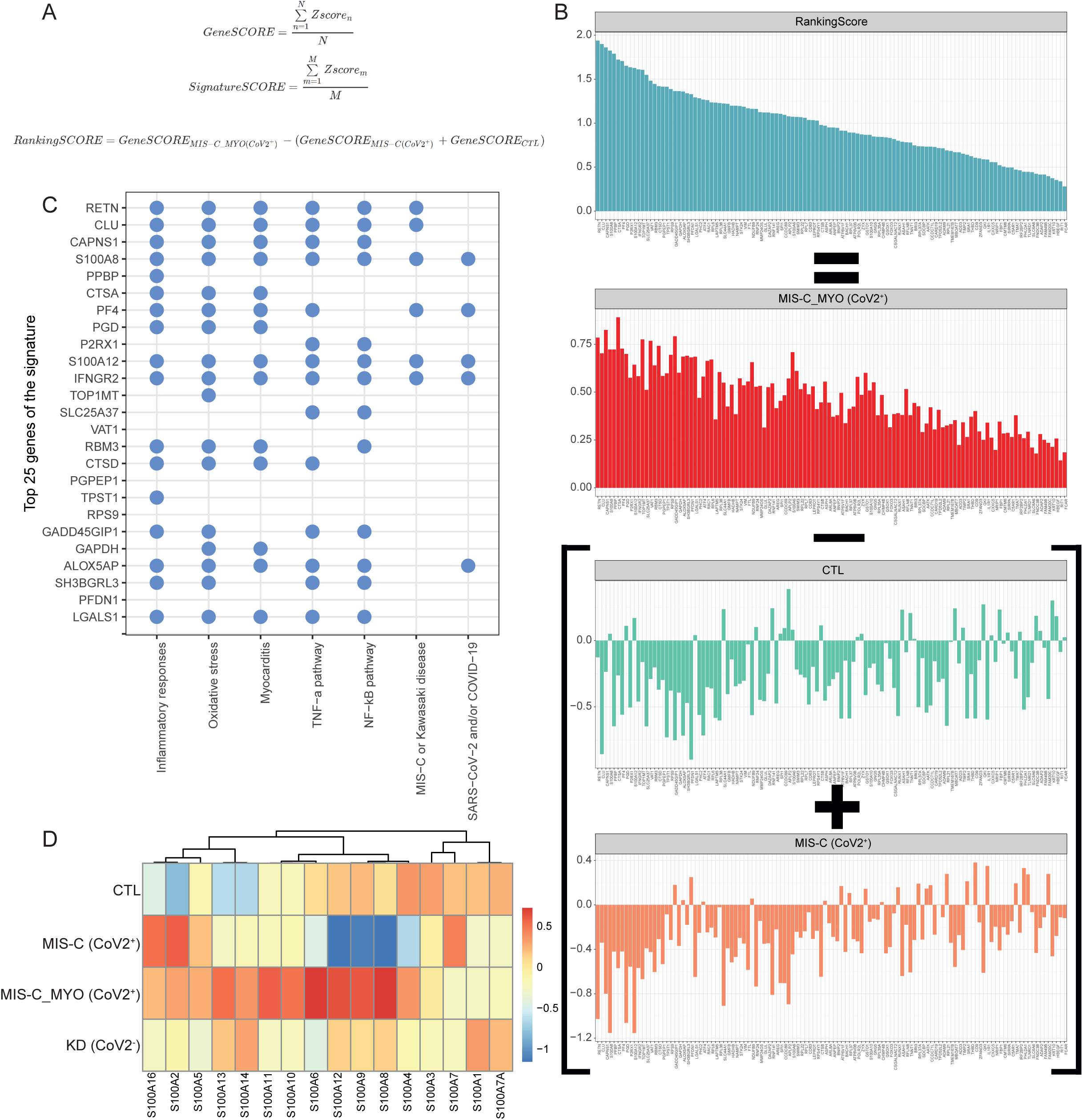
SignatureScore and top genes correlated with the occurrence of myocarditis in MIS-C (CoV2+). Related to Figure 7. **A.** *MIS-C_MYO (CoV2^+^)*-specific genes were scored (GeneSCORE) by adding their expression values (as z-scores) in each sample, divided by the total number of samples. Similarly, a SignatureSCORE was derived for each sample as the mean z-score of these samples. From these two scores, a RankingScore was calculated by subtracting the GeneScore of the *MIS-C (CoV2^+^)* and the CTL groups to the GeneSCORE of the *MIS-C_MYO (CoV2^+^)* group. **B.** Top: Barplot of the RankingScore of each gene. Red, green and orange barplots: GeneScore of the *MIS-C_MYO (CoV2^+^)*-specific genes in the *MIS-C_MYO (CoV2^+^)*, CTL and *MIS-C (CoV2^+^)* groups respectively. Genes are ranked based on their RankingScore as explained in S7A **C.** Curated analyses based on literature regarding the known links between the top 25 genes, and inflammatory response, myocarditis, oxidative stress, TNF-α, NF-κB, MIS-C or Kawasaki disease, SARS-CoV-2 and/or COVID-19. On the Y axis, genes are positioned from top to bottom following their rank (Top = ranked 1). A blue dot means strong links between the gene and the pathway (on the X axis) were found in peer-reviewed scientific publications. **D.** Heatmap of the expression of the genes of the S100A family, extracted from Bulk-RNA-SEQ, from PBMCs. Color scale indicate the scaled GeneSCORE (mean z-score of the gene in all samples of a group), with red and blue representing the highest and lowest expressions respectively. Hierarchical clustering of the genes was computed with a Pearson’s correlation as a distance.

**Figure S8.**
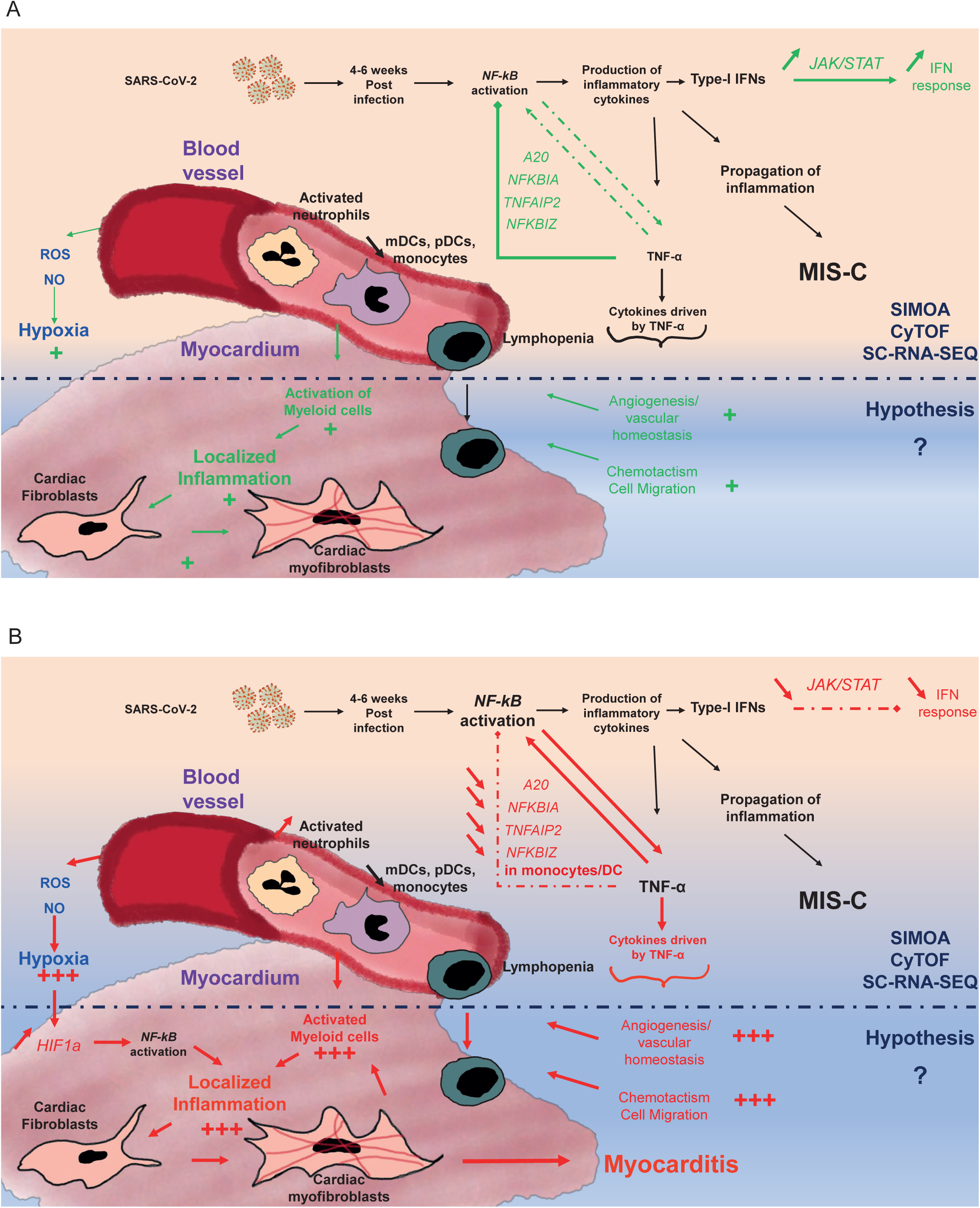
**A & B.** Graphical representation based on cytokines, cellular and transcriptomic analyses (part above black dotted line), combined with known literature (part under the black dotted line), illustrating a putative model explaining the occurrence of myocarditis among children in the *MIS-C (CoV2^+^)* group. **A.** model based on the *MIS-C (CoV2^+^)* group and **B.** model based on the *MIS-C_MYO (CoV2^+^)* group. Black writing represents genes and functions both modulated in the *MIS-C (CoV2^+^)* and *MIS-C_MYO (CoV2^+^)* groups compared to CTL, whereas green and red are highlighting genes and pathways differentially modulated in the *MIS-C (CoV2^+^)* and *MIS-C_MYO (CoV2^+^)* groups, respectively.

